# Decoding Collagen’s Thermally Induced Unfolding and Refolding Pathways

**DOI:** 10.1101/2024.10.02.616390

**Authors:** Alaa Al-Shaer, Nancy R. Forde

## Abstract

Collagen has been evolutionarily selected as the preferred building block of extracellular structures. Despite inherent thermal instability of individual proteins at body temperature, collagen manages to assemble into higher-order structures that provide mechanical support to tissues. Sequence features that enhance collagen stability have been deduced largely from studies of collagen-mimetic peptides, as the large sizes of collagens have precluded high-resolution studies of their structure. Thus, there is a need for new methods to analyze the structure and mechanics of native collagen proteins. In this study, we used AFM imaging to investigate the temperature response of collagen types I, III and particularly IV. We observed a time-dependent loss of folded structures upon exposure to body temperature, with structural destabilization along the collagenous domain reflected by a shorter overall contour length. We characterized the sequence-dependent bending stiffness profile of collagen IV as a function of temperature and identified a putative initiation site for thermally induced unfolding. Interchain disulfide bonds in collagen IV were shown to enhance thermal stability and serve as the primary nucleation sites for *in vitro* refolding. In contrast to the canonical C-to-N terminal folding direction, we found an interchain cystine knot to enable folding in the opposite direction. Multiple sequence alignments revealed that this cystine knot is evolutionarily conserved across metazoan phyla, highlighting its significance in the stabilization of early collagen IV structures. Our findings provide mechanistic insights into the unfolding and refolding pathway of collagen IV, providing valuable insight into how its heterogeneous sequence influences stability and mechanics.

## Introduction

Collagens emerged at the dawn of metazoan evolution, introducing extracellular proteins pivotal for the development of multicellular life.^1^ One of the many physiological roles for this class of proteins is as an essential component of basement membranes (BMs), specialized sheet-like extracellular matrices that support many tissues and guide their morphogenesis.^2,3^ BMs serve as an interactive scaffold that can mechanotransduce information to regulate cellular behavior.^4^ Among the key determinants of BM mechanical properties is the network-forming collagen type IV. Genomic analyses have traced the origins of canonical collagen IV genes to early metazoan ancestors,^1,3,4^ implying that collagen IV played a vital role in the expansion and diversification of metazoan life.^1,5^

The most abundant and ubiquitous isoform of collagen IV is the (α1)_2_α2 heterotrimer, which is consistently found in nearly all BMs across Metazoa.^6^ Collagen IV synthesis begins in the endoplasmic reticulum of the cell, where three left-handed polyproline type-II α-chains fold into a right-handed triple helix to form the collagenous domain of the protein. The C-terminal non-collagenous (NC1) domain serves as the recognition site that mediates association of the α-chains and nucleates the triple helix folding, which proceeds from the C to the N terminus in a zipper-like fashion.^7^ The contour length of the collagenous domain is approximately 360 ± 20 nm, which includes an N-terminal triple-helical 7S domain.^8^ The folded protein structure, containing the NC1 and collagenous domains, is referred to as a “protomer”. Distinct from fibrillar collagen types, the collagenous domain of collagen IV is not only significantly longer, but also contains structural heterogeneities owing to the presence of interruptions in the triple-helix-defining (G–X–Y)_n_ repeating tripeptide unit in each α-chain (where X and Y are often P = proline and O = hydroxyproline). These interruptions, which lack glycine as every third amino acid, enhance the mechanical flexibility of the protomer, but may also delay triple helix folding.^8–11^ Folding delays have been observed in collagen-mimetic peptides and in non-natural, disease-associated interruptions of the wildtype collagenous G-X-Y repeat in other collagens,^10,11^ but the role of endogenous interruptions in the folding pathway of collagen IV folding is less clear.^12,13^

Considering collagen’s pivotal role in assembling into higher-order structures that provide mechanical support to surrounding tissues, it is both fascinating and perplexing that the protein is thermally labile at body temperature.^14^ Nature fine-tunes the melting temperature of collagen in different species to be just below, rather than above, body temperature, primarily through adjustments in hydroxyproline content.^15,16^ The G–P–O repeating unit, commonly found in natural collagens, represents the most thermally stable triple-helix-forming triplet with a melting temperature in peptide studies well above body temperature (e.g. *T*_m_ = 47°C).^17^ However, natural collagen sequences are heterogeneous, resulting in their significantly lower melting temperatures.^23–25^ The sequence heterogeneity of collagen leads to variable mechanics in different regions of the protein, and is vital for upholding its biological function, despite contributing to its thermal instability.^8,18–20^ While hydroxyproline is a key promoter of collagen stability, other features such as salt bridges and interchain disulfide bonds can also contribute.^21,22^ Collagen’s high frequency of charged residues has been linked to improved folding efficiency and triple helix stability through salt bridge formation.^23,24^ As collagen folds and adopts its native state, its structure can be stabilized by the formation of interchain disulfide bonds in the oxidizing environment of the ER.^13^

Not only is collagen’s melting temperature below body temperature, but its denaturation is essentially irreversible on practical timescales.^13,14,25,26^ The “agonizingly” slow refolding dynamics (on the order of hours) arise in part from the repetitive G-X-Y sequence,^31^ which can refold into incorrectly matched yet still locally triple-helical structures, whose annealing dynamics depend on the local sequence.^7,25^ In collagen IV, chain misalignment is likely to be exacerbated by the G-X-Y interruptions. It is plausible that disulfides found within the collagenous domain of collagen IV can act as structural clamps and guide refolding, as they do for collagen III.^7,26,27^ The existence of such structural clamps could help to explain how collagen IV retains its structure in spite of its length and sequence complexity.

Conventional methods used to investigate collagen structure and stability have provided valuable insights, but they come with limitations. Collagen model peptides, while suitable for crystallization and NMR studies, cannot capture the inherent sequence heterogeneity within natural collagens. Additionally, bulk assays such as circular dichroism (CD) and differential scanning calorimetry (DSC) primarily characterize ensemble properties, and thus cannot elucidate folding/unfolding pathways followed by individual proteins within the ensemble. The occurrence of rare and transient events like micro-unfolding (localized breathing of the triple helix) have been inferred from bulk assays,^14,20^ yet have not been directly observed. Thus, new methods areneeded to establish correlations between collagen’s sequence, structure, and mechanics within the context of the full-length protein.

Atomic Force Microscopy (AFM) provides a promising avenue for studying collagen at high spatial resolution.^8,28–30^ In this study, we use AFM to explore how temperature influences the structure and mechanics of collagen IV protomers. Our work provides mechanistic insights into the transition of collagen IV from its folded triple-helical state to a collapsed random coil, and its refolding. We compare these findings to those in the less structurally complex fibrillar collagens (I and III) that lack triple-helical interruptions. Additionally, we investigate the role of disulfide bonds in contributing to the thermal stability and reversibility of collagen denaturation. Among the probed disulfide bonds, we highlight the functional significance of an evolutionarily conserved cystine knot in collagen IV.^31^ Through our investigation, we elucidate roles of specific structural elements in collagen IV and propose mechanisms for its unfolding and refolding pathways.

## Results

### Characterization of Purified Collagen IV

Collagen IV with an (α1)_2_α2 composition was obtained from the cultured media of the murine epithelial cell line PFHR9. The protein was purified by adapting a previously described collagen IV protomer purification protocol (methods).^13,32^ A representative AFM image of the purified full-length collagen IV protomer, with a globular C-terminal NC1 domain and a collagenous triple-helical domain, is shown in Fig. 1A. The triple-helical quality of the sample was measured by CD spectroscopy. The spectra exhibited a maximum at 222 nm and a minimum at 198 nm, with an *R*_pn_ value of 0.13 (see methods), indicating a triple-helical structure (Fig. S1).

**Figure 1.**
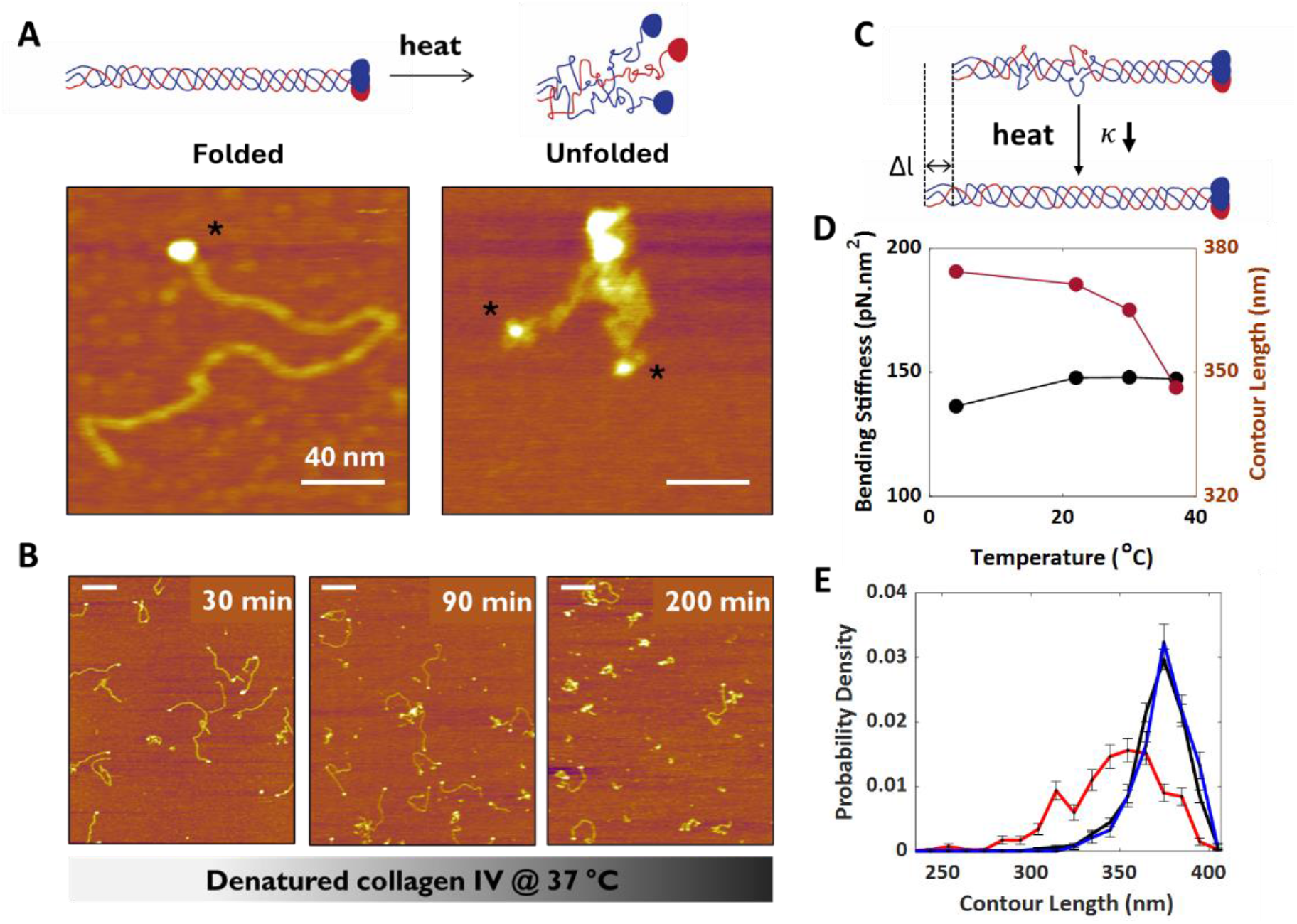
Collagen IV characterization and response to temperature. **A**. Structural representation (top) and corresponding AFM image (bottom) of collagen IV in its folded state (left) compared to an unfolded collapsed state (right). The NC1 domains are denoted with an asterisk, indicating their association into a trimer in the folded state and collapse into monomers in the random coil. Scale bars = 40 nm. **B**. AFM images illustrating the time-dependent denaturation of collagen IV upon heating at 37°C. Scale bars = 200 nm. **C**. Proposed effect of temperature on the contour length and bending stiffness κ of collagen IV below its melting point. **D**. Bending stiffness and contour length of collagen IV versus temperature after 30 min incubation periods. **E**. Distribution of contour lengths of collagen IV with visibly intact NC1 domains, at different temperatures.

### Thermal Denaturation of Collagen IV

Collagen IV was incubated in PBS (pH 7.0) at various temperatures, and we observed a decrease in folded structures above 37°C (Figs. 1C, S2). A structure was classified as “folded” if the protomer possessed a near-native length and an intact NC1 trimer cap. At 37°C, we imaged collagen IV after different incubation periods and observed its time-dependent denaturation (Figs. 1C, S3). This denaturation process involves the separation of α-chains including the dissociation of the NC1 trimer into monomers, which can be seen within a collapsed structure (Fig. 1A). Consequently, the energetically preferred conformation of the collagen IV protomer in a physiologically relevant environment is an unfolded, collapsed structure rather than a triple helix.^14,33^ Comparably, we note similar destabilization in the uninterrupted fibrillar collagen types, namely collagen I and III, with an even more pronounced destabilization in an acidic environment (Fig. S2).

We sought to determine how collagen IV transitions from its native triple-helical state to a random coil. We assessed its conformation at different temperatures by tracing chains from AFM images, by first focusing on those with visually intact collagenous and NC1 trimer domains, which we consider “folded”. This approach was aimed at providing a detailed examination of the *initial* internal structural changes within chains that had not yet unfolded into a collapsed state. We hypothesized that temperature would affect collagen’s mechanics by inducing the formation of local breathable regions along its contour. Additionally, we hypothesized that these micro-unfolded regions would contract, transforming into local flexible hinges, and lead to a reduction in both contour length and bending stiffness *κ* (Fig. 1D). Such thermally induced softening has been observed in DNA, where a decrease in bending stiffness has been attributed to local breathing of the double helix.^34^ We tested this hypothesis for collagen IV by determining the contour lengths of the folded proteins from traced AFM images and by analyzing the shapes of these contours to determine the global bending stiffness using the worm-like chain model (equations 1, 2, 4 in methods).

Surprisingly, folded collagen IV protomers showed no significant decrease in global bending stiffness as temperature increased from room temperature (22°C; Fig. 1E). This observation was inconsistent with our hypothesis, in which micro-unfolding bubbles nucleating throughout the collagenous domain would manifest in a softening of bending stiffness at physiological temperatures. Interestingly, collagen IV appeared to be less stiff at 4°C, which may be due to imperfections in the triple helix arising from disruptions in intermolecular hydrogen bonding networks at lower temperatures rather than thermally activated micro-unfolding.^35^ Although the folded chains that we traced showed no significant mechanical changes at elevated temperatures, they did exhibit a structural change consistent with our micro-unfolding hypothesis: a decrease in contour length (Fig. 1E). This decrease in average contour length is accompanied by a broadening of the contour length distribution compared with ambient conditions (at 37°C, ⟨*L*⟩ = 346 nm, σ = 27 nm; at room temperature, ⟨*L*⟩ = 371 nm, σ = 16 nm; mean and standard deviation) as seen in Fig. 1E. This increased distribution of lengths of the collagenous domain demonstrates a heterogeneity of response to temperature, the origins of which we aimed to elucidate.

To determine whether particular regions within collagen IV are more prone to local unfolding than others, we performed a position-dependent analysis of the local bending stiffness (equations 3, 4),^8^ again focusing on chains with an intact NC1 domain. This analysis provides a measure of the effective bending stiffness *κ*(*s*) over 30 nm segments centered at nanometer increments of *s* along the length of the collagenous domain. In Fig. 2A, an amino acid representation of the full sequence of mouse (α1)_2_α2 collagen IV is presented and aligned with the calculated effective bending stiffness profiles in Fig. 2B. In our previous study, we observed variable mechanics along the length of tissue-derived collagen IV, with increased flexibility in the N-terminal quarter correlating with a higher extent of interruptions.^8^ Peaks in stiffness aligned with long triple-helical stretches, and minima aligned with interruptions in all three α-chains or overlapping interruptions. Our current finding indicates that PFHR9-derived collagen IV exhibits a comparable mechanical profile to this previously characterized, tissue-derived collagen IV (Fig. S4).

**Figure 2.**
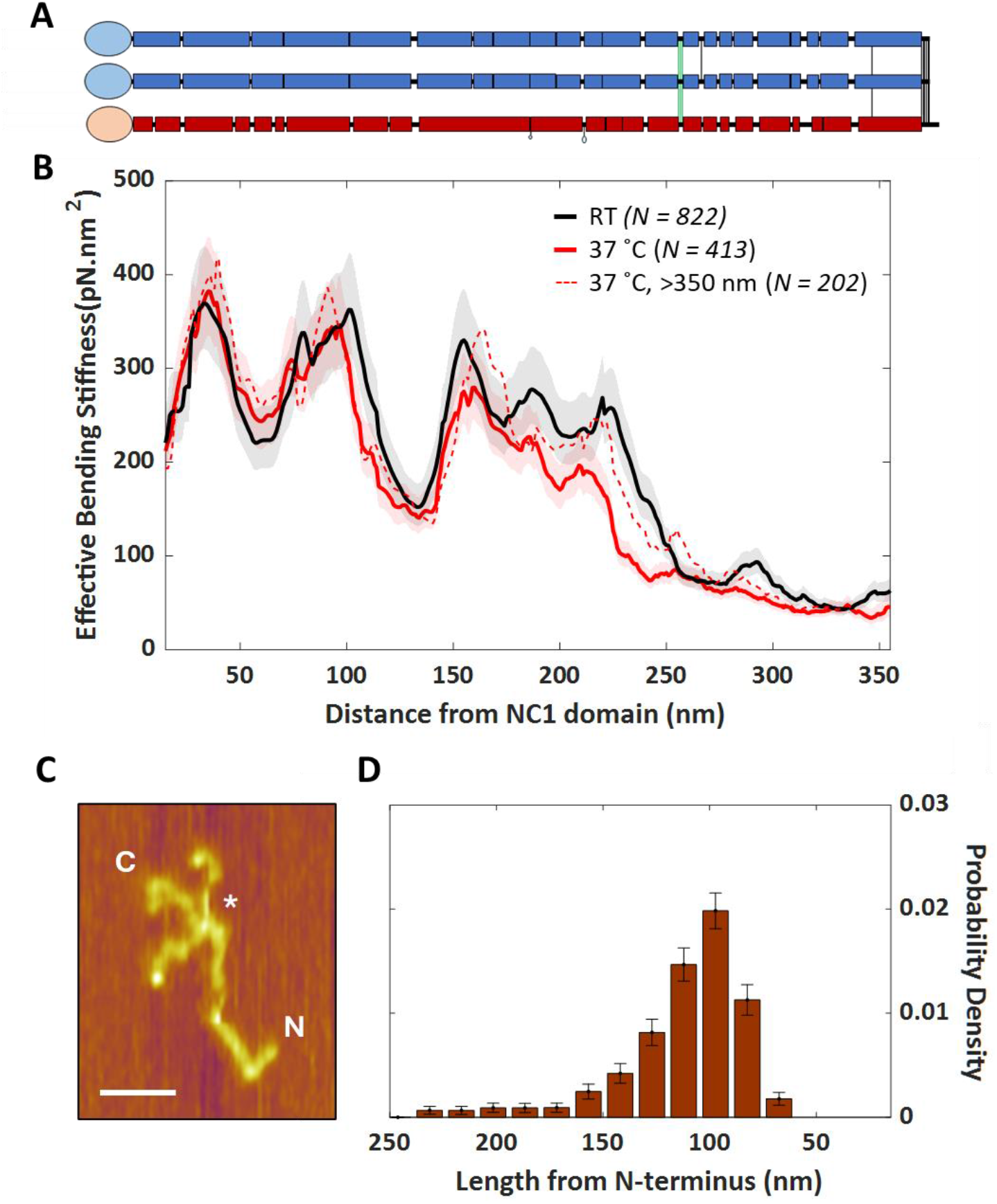
Collagen IV bending stiffness profile at different temperatures, and evidence for a partially unfolded intermediate. **A**. Schematic representation of the α1 and α2 amino acid sequences from mouse collagen IV. Boxes indicate sequence regions containing (GXY)_n_, while horizontal black lines indicate interruptions in this pattern. Vertical lines between the chains identify potential interchain disulfide bonds, with green denoting the cystine knot. **B**. Effective bending stiffness profile of collagen IV deposited at room temperature (black) or after a 30-minute incubation at 37°C (all chains: red; contour length >350 nm: dashed red). **C**. An AFM image of a collapsed structure showing a retained triple-helical segment at the N-terminal end. Scale bar = 50 nm. **D**. Length distribution of N-terminal intact domains (*N* = 300). Intermediate structures were traced starting from the N terminus to the point at which fraying was observed (* in panel C). *x* axes in panels B and D are aligned by position along the collagen IV chains. The most probable and median N-terminal folded lengths (105 nm) coincide with the position of the cystine knot.

We then compared the effective bending stiffness profile of folded collagen IV protomers at 37°C with their profile at room temperature (Fig. 2B). Overall, these profiles were very similar, indicating no significant structural softening at 37°C, as seen already by the comparison of global bending stiffness (Fig. 1E). However, a distinct response at 37°C was found in a region 200-250 nm from the NC1 domain. We sought to determine whether chains exhibiting a decrease in contour length contributed to this localized softening and apparent destabilization. We found that the bending stiffness profiles of shorter (length < 350 nm) and longer (>350 nm) chains differed significantly only in the region 200-250 nm from the NC1 domain (Fig. S5). Longer chains possessed the same bending stiffness in this region as collagen IV at room temperature, implying no destabilization (Fig. 2B), whereas the shorter chains exhibited increased flexibility in this region (Fig. S5). These findings support our hypothesis that increased temperature induces structural destabilization accompanied by shortening. However, we note that destabilization seems to be localized to this one region of collagen IV. The destabilization observed in this region hints at a potential nucleation site for complete unfolding of collagen IV. Below, we refer to these shorter protomers with increased flexibility in the 200-250 nm region as “unfolding-prone” structures. Of the ∼2/3 of protomers that remained folded after 30 minutes of incubation at 37°C, approximately half are associated with this unfolding-prone state while the other half remain fully structured.

Conversely, when examining the structures that collapsed upon heating, *i*.*e*., those in which the NC1 trimer was not intact, we identified a subset that remained partially folded and retained an intact N-terminal collagenous segment. While there is variability in the length of these segments, there is a clearly preferred value of ∼105 nm (*N*=300; Fig. 2D). Examining the sequence of collagen IV near this region, we find two cysteine residues in each of the three α chains, which are one (in α2) or two (in α1) residues apart. These are the first cysteines encountered when unfolding the collagenous domain from the NC1 C terminus. The six cysteine residues in this region can form three disulfide bonds to crosslink the three α-chains, creating a so-called cystine knot (interchain disulfides denoted with green lines, Fig. 2D).^36^ This cystine knot, along with other interchain disulfide bonds within the 7S domain, anchor the three α-chains together and likely contribute to the stability of this retained segment of the collagenous domain.

These sequence-dependent observations lead us to infer that there is a cooperative unfolding block in collagen IV, spanning the region from the NC1 domain to the cystine knot. We reach this conclusion because (1) when NC1 trimers are visibly disrupted, there are very few collagenous domains with lengths longer than the N-terminal cystine-linked segment; and (2) when NC1 trimers are intact, the only disruption in the collagenous domain that we detect corresponds to localized breathing in the region immediately C-terminal to the cystine knot. Our findings thus suggest that micro-unfolding occurs C-terminal to the cystine knot, and unfolding propagates from this point rapidly and apparently in a highly cooperative fashion towards the NC1 domain.

### Refolding of Collagen IV

Given that collagens can fold in a zipper-like fashion intracellularly without the aid of folding chaperones,^7, 13, 26^ we next assessed how collagen IV refolds *in vitro*.

We first determined whether the destabilized region detected in the bending stiffness profile of collagen IV at 37°C can be restored by cooling. By examining those collagens that appear folded, as we did at 37°C, we found that the destabilized region returned to its original stiffness upon cooling, whether to room temperature or to 4°C (Fig. S6A). This result highlights the ability of collagen IV to refold and regain local structure. Furthermore, it implies that destabilization of this region is thermally reversible, with dynamics that could be described as micro-unfolding or breathing of the triple helix. We also note that the contour lengths of protomers are returned to their native distribution upon cooling (Fig. S6B).

Having established this reversible *local* unfolding within collagen IV, we next explored whether collagens that had unfolded to a collapsed state (Fig. 3A) could fold back to their native state. To ensure a starting population of collapsed structures, collagen IV was heated to 43°C for 30 minutes. After subsequent overnight cooling at room temperature, we found that collagen IV could refold, with the refolded proteins (314/466 structures) visually resembling the initial sample (Fig. 3B). The other proteins were found to adopt non-native structures, many with NC1 domains that had not reformed trimers (of the type shown in Fig. 3C).

**Figure 3.**
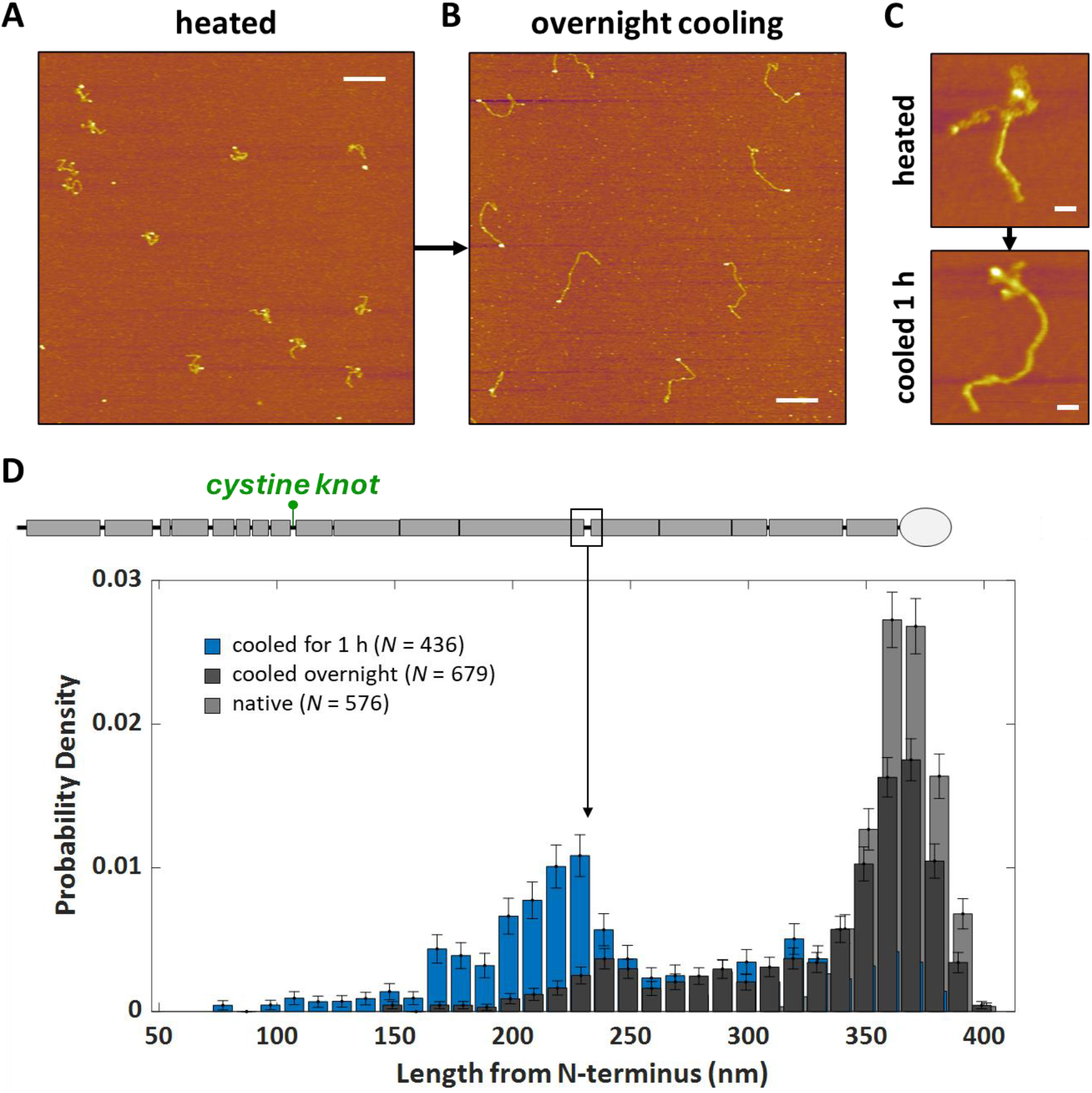
Collagen IV autonomously refolds towards the C terminus upon cooling. **A**. AFM image of collagen IV after heating at 40°C for 90 minutes. Scale bar = 200 nm. **B**. AFM image of collagen IV following overnight cooling at room temperature. Some incomplete refolding is evident, e.g. protein in lower left. Scale bar = 200 nm. **C**. AFM image of a collapsed collagen IV structure after heating at 40°C, showing a retained N-terminal triple-helical segment (top). AFM image of a partially refolded collagen IV structure after a subsequent 1 hour of cooling at room temperature (bottom). The length of the triple-helical segment in partially folded structures lengthens after cooling. Scale bars = 50 nm. **D**. Distribution of contour lengths, measured from the N terminus. Shown are distributions for samples cooled for 1 h, cooled overnight, and from native collagen IV. The increase in length indicates refolding towards the C terminus. The location of the largest overlapping interruption is denoted with an arrow.

We then investigated how the presence of interruptions influences the structures observed in the refolding process. While repeating units of G–X–Y form a triple-helical structure, interruptions in this pattern introduce an entropic barrier, likely delaying the zippering of the remaining length of α-chains until triple helix renucleation occurs beyond the interrupted site. Previous studies confirm the slower folding progression of mutant collagens from diseased patients and of recombinant collagens containing interruptions.^10,11^ Here, we first examined refolding after only one hour at room temperature. At this time point, very few of the structures have refolded to their native length. Instead, many have collagenous domains extending up to only 240 nm from the N terminus (Fig. 3D). This implies that they have undergone partial refolding past the location of the cystine knot (110 nm from the N-terminus) but are held up from further refolding by a free energy barrier. By aligning the contour lengths from the N-terminus with the amino acid representation of collagen IV, we find that the largest overlapping interruption is located 240 nm from the N-terminus, where we see the largest population of intermediate collagenous domain lengths (arrow, Fig. 3C). This provides direct evidence that natural interruptions act as folding barriers in the wild-type protein, likely entropic in origin.^37^ Intriguingly, these results indicate N-to-C directional folding of the collagenous domain *in vitro* (Fig. 3D), contrary to the natural folding direction of collagens within the cell.^7^

While samples cooled for many hours predominantly contained structures that had refolded back to native length (Fig. 4A), several structures appeared misfolded within the triple helix or at the NC1 domain (Fig. 4B). A bleb was evident in some structures that also displayed a commensurately shorter contour length, indicating a bulge and lack of proper refolding within the triple helix. To examine whether the refolded collagen IV population displays comparable mechanics to the native structure, we traced all structures that emerged after overnight cooling, focusing on those with a distinct NC1 trimer and no evident unstructured regions within the collagenous domain. While the effective bending stiffness did not exactly match the native collagen IV profile at ambient temperature, it did exhibit qualitative similarities in maxima and minima (Fig. S7A). When we narrowed our analysis to chains with contour lengths above 350 nm (comparable to the full-length, natively folded protein), the resulting effective bending stiffness profile closely resembled that of native collagen IV (Fig. 4C). By contrast, when limiting our analysis to chains shorter than 350 nm, we failed to capture the characteristic local maxima and minima of collagen IV’s mechanical signature (Fig. S7B). This suggests that visual appearance alone cannot be used to classify collagen IV as refolded, and that within this population, there are shorter chains, corresponding to incompletely folded structures that exhibit distinct bending stiffness profiles. We note that, even for the longer contour length subset of chains, the C-terminal region of the refolded chains (between the NC1 domain and the overlapping interruption) is more flexible on average than the native structure (Fig. 4C). This suggests that even this population of near-native length proteins contains some misfolding, leading to a reduction in triple-helical content and hence increased local flexibility.^8^ The refolding challenges faced by collagen IV, particularly from the interruptions, have been previously noted.^12^ These observations highlight the critical role of proper folding in restoring the structural integrity and mechanical characteristics of collagen IV.

**Figure 4.**
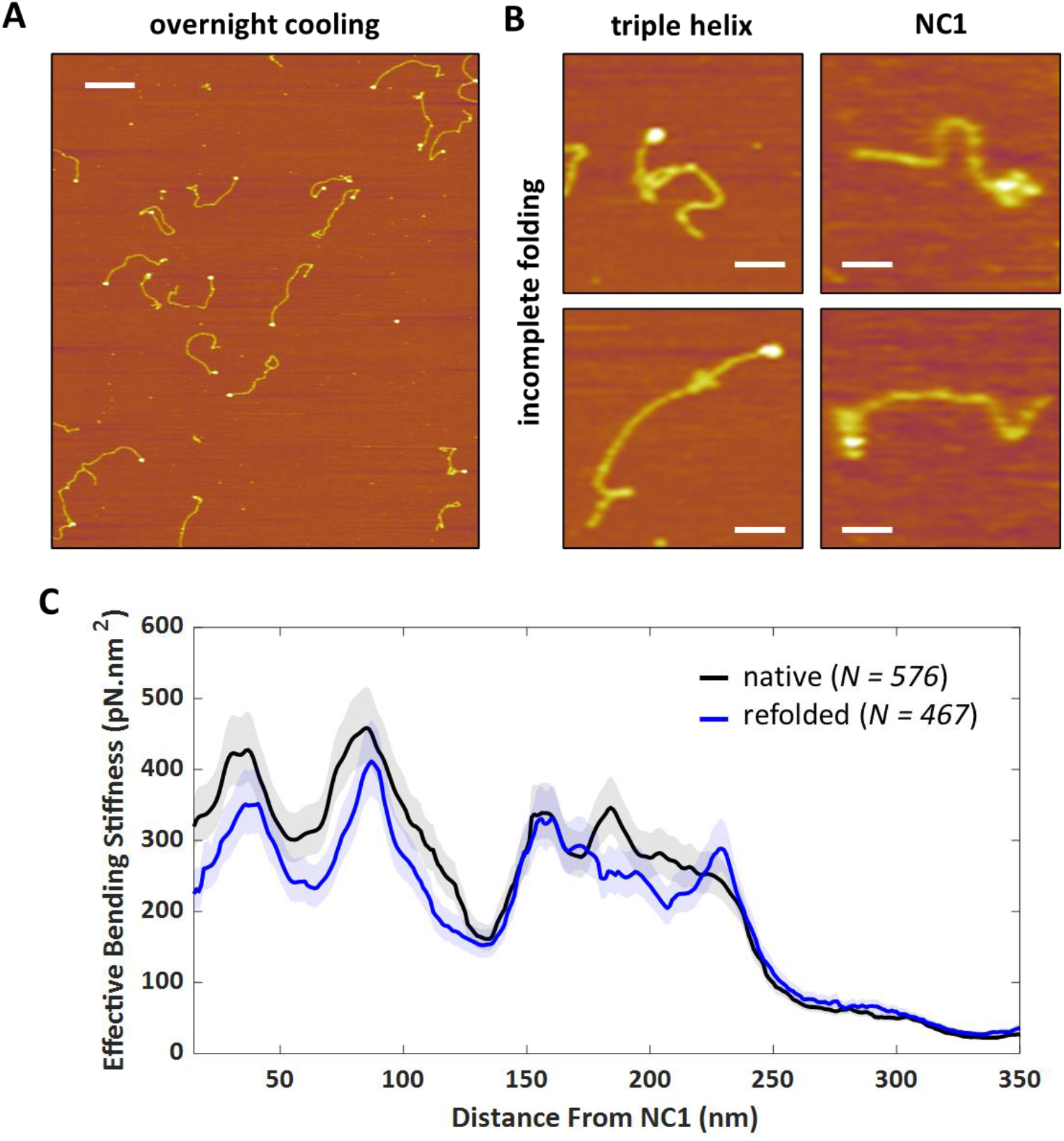
Mechanics of refolded collagen IV. **A**. AFM image of collagen IV heated at 43°C followed by overnight cooling at room temperature. Scale bar = 200 nm. **B**. Example images of incompletely folded structures within the triple helix (e.g. blebs) or at the NC1 domain. Scale bars = 50 nm. **C**. Effective bending stiffness profile at room temperature of refolded collagen IV for chains with a contour length >350 nm (blue), and native collagen (black).

### Denaturation and Refolding of Fibrillar Collagens

Given the structural complexity of collagen IV, particularly due to interruptions within its collagenous domain, we wished to compare its temperature response with that of the less structurally complex fibrillar collagens. As seen previously, fibrillar collagens possess a significantly greater (2x) bending stiffness than collagen IV (Fig. S8),^8,28^ due to interruptions in the latter.^8,13,38^ Similar to what we observed for collagen IV, intact collagens I and III also displayed no change in bending stiffness between room temperature and 37°C (Fig. S8), and also decreased in contour length at this elevated temperature (Figs. S9, S10). On rare occasions, we observed collagen end-fraying (arrow, Fig. S9A), shedding light on a possible shortening mechanism of uninterrupted collagens. We return to this point in the Discussion section.

The role of interchain disulfide bonds in nucleating refolding in collagens has been reported, particularly for collagen III.^25^ Our findings confirm that collagen III, containing a cystine knot at its C terminus, exhibited refolding capability, while collagen I, devoid of cysteines, failed to refold upon cooling (Fig. S9C). Given the essential role of interchain disulfide bonds in facilitating refolding of fibrillar collagens, we sought to determine whether this mechanism holds true for collagen IV.

### Reduction of Collagen IV

To test whether disulfide bonds act as nucleation sites for refolding of collagen IV, we reduced the cysteines (Fig. 5A), heated, cooled, and imaged the resulting collagen IV structures. Unlike the previously observed collapsed collagen IV structures, here we predominantly observed structures that resemble α-chain monomers (Fig. 5B). These structures were more compact and thinner, with no examples of denatured protomers displaying three NC1 domains in proximity or appearing to retain an intact triple-helical segment. Instead, in the absence of interchain disulfides, the α-chains disperse randomly in solution upon denaturation. In further contrast to the heated, non-reduced collagen, here we also observed aggregates which we speculate may be induced by interactions between NC1 domains. While the melting temperature of NC1 trimers is significantly higher (*T*_m_ ≈ 66°C) than the temperatures used in this study, reduction of disulfides within each domain may reduce thermal stability, leading to aggregation.^39^ Following overnight cooling at room temperature, no native-like collagen IV structures emerged (Fig. 5B), confirming the role of disulfides in nucleating refolding.

**Figure 5.**
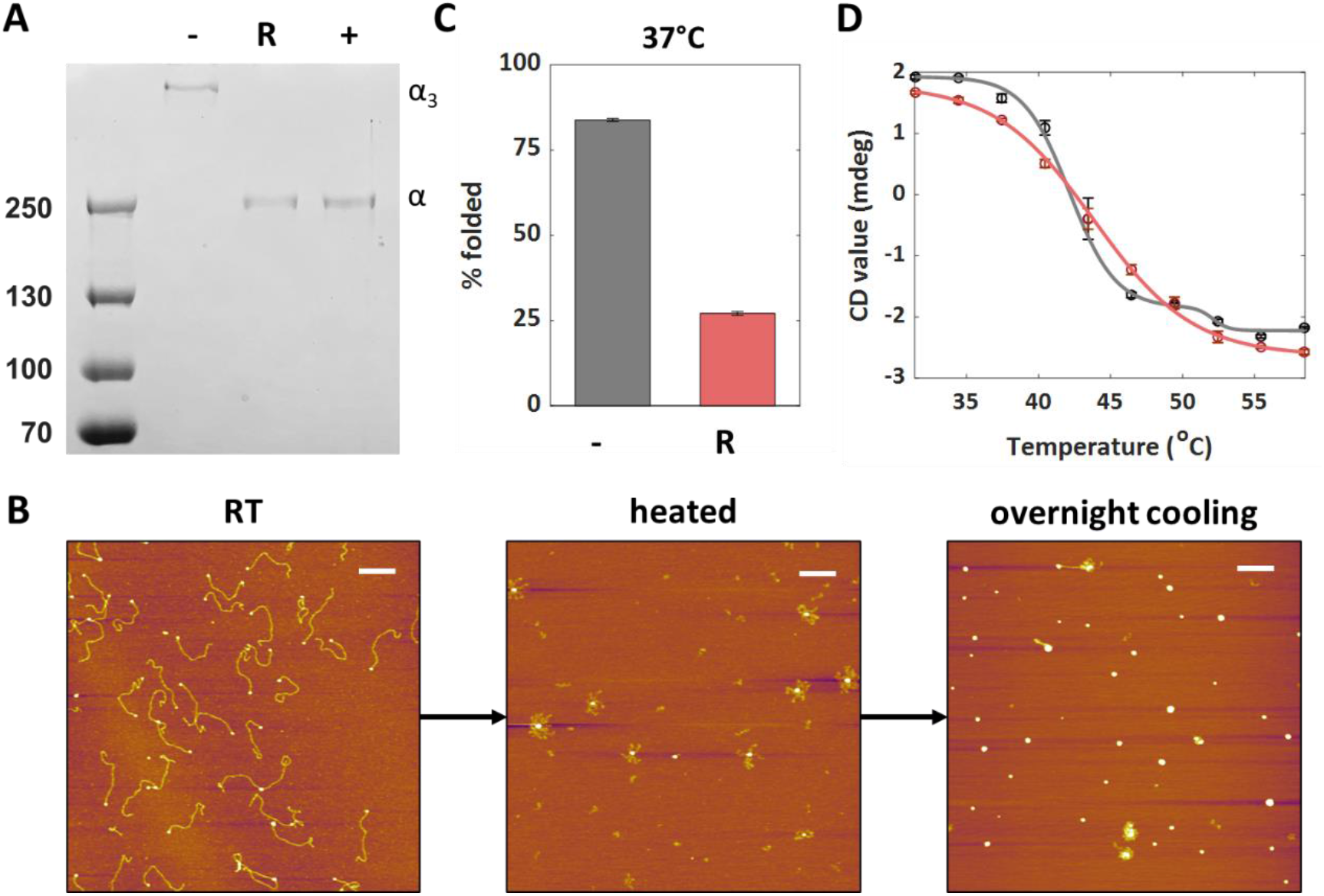
Disulfide bonds contribute to thermal stability and serve as nucleation sites for refolding. **A**. SDS-PAGE of collagen IV run with non-reducing loading dye (negative control, -), after treatment with TCEP-HCl and run with non-reducing loading dye (R) and run with reducing loading dye (positive control, +). α indicates the bands associated with single α1 and α2 chains; α_3_ indicates the location of three crosslinked α chains. **B**. AFM images of reduced collagen IV imaged at room temperature, after heating at 40°C, and following overnight cooling at room temperature. No refolding was observed. Scale bars = 200 nm. **C**. Comparison of the percentage of folded structures after heating non-reduced (−) and reduced (R) collagen IV at 37°C for 30 minutes. **D**. Thermal denaturation profiles obtained by monitoring the CD signal at 222 nm of reduced collagen IV (red; monophasic) and non-reduced collagen IV (gray; biphasic).

Next, we investigated whether the disulfide bonds in collagen IV contribute to the stability of the folded structure. We found that reduced collagen IV, incubated at 37°C for 30 minutes, displayed more unfolded structures than non-reduced collagen IV, implying reduced thermal stability upon chemical reduction (Fig. 5C, S11). (Here and elsewhere, thermal stability denotes kinetic stability in response to temperature changes.^40^ To obtain a thermal denaturation profile for collagen IV, we measured a melting curve with CD spectroscopy, monitoring the intensity at 222 nm as the sample was heated from 30 to 60°C at a rate of 0.5°C/min (Fig. 5D). These measurements revealed that reduced collagen IV began unfolding earlier and followed a monophasic transition with a melting temperature of 42.4 ± 0.2 °C. Conversely, the melting profile of non-reduced collagen IV was not well described by a monophasic model, but instead exhibited a multiphasic transition. A biphasic model provided a relatively good fit to the data under both neutral and acidic pH conditions (Fig. 5D, Fig. S12), consistent with prior research.^13^ The steepness of the first and dominant denaturation transition for non-reduced collagen IV implies that it is cooperative (Fig. 5D). Comparing with our AFM results, we propose that this represents the cooperative unfolding of the block C-terminal to the cystine knot to the NC1 domain. The second and smaller transition in the CD melt curve corresponds to the unfolding of a more thermally stable and shorter triple-helical segment. Its absence in the denaturation profile of reduced collagen IV suggests that this segment, which requires a higher temperature for complete unfolding, corresponds to the N-terminal triple-helical segment observed in our AFM images, which is stabilized by multiple interchain disulfide bonds (Fig. 2A). This attribution is consistent with the higher thermal stability previously observed for the N-terminal 7S domain of collagen IV.^41^ Overall, our findings demonstrate that disulfide bonds not only anchor the α-chains together and maintain registry alignment, but can serve as a nucleation site for refolding, and endow thermal stability.

### Cystine Knot across Metazoa

Given collagen IV’s presence across Metazoa,^1,42^ we investigated the occurrence and significance of the cystine knot across various species spanning major animal phyla (Fig. 6). Previous work has found this structural element to be conserved in four distant organisms including fruit flies, roundworms, and Cnidaria.^31^ Expanding on this study, we performed a multiple sequence alignment of COL4 sequences across 14 species from 9 major phyla (Table 1) and found a conserved pattern of two cysteine residues, always separated by two residues. Although this pattern was primarily analyzed in the α1 chain and its subfamily, we refer to it as a cystine knot because it is also observed in all isoforms (cysteines in all three α-chains can establish the cystine knot^22^), including α2, α3, α4, α5, and α6 (from *M. musculus*, data not shown). Interestingly, in α2 and α6 chains, the two cysteines are separated by just one residue.

**Table 1.**
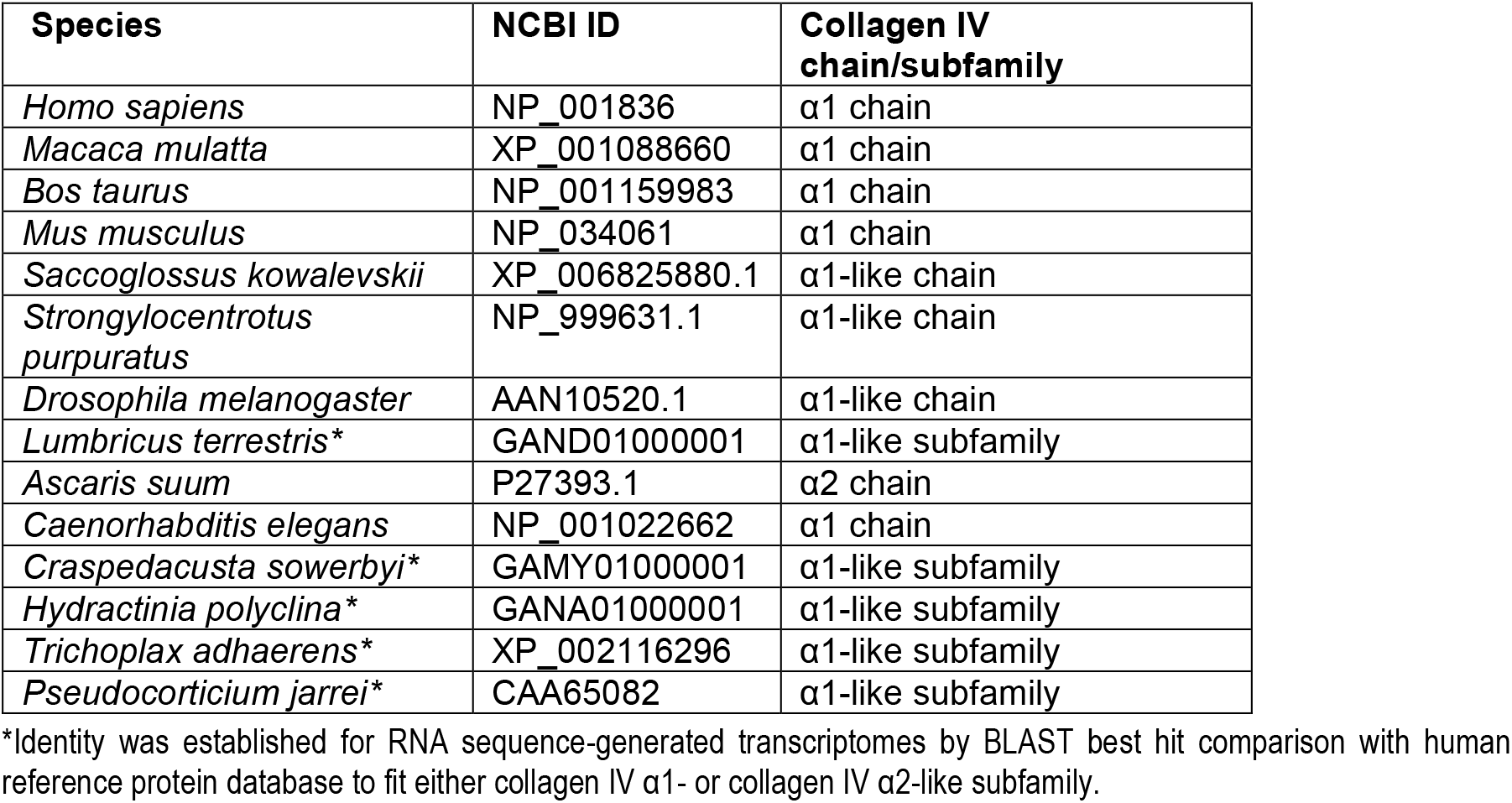
National Center for Biotechnology Information Identification accession numbers for collagen IV sequences.

**Figure 6.**
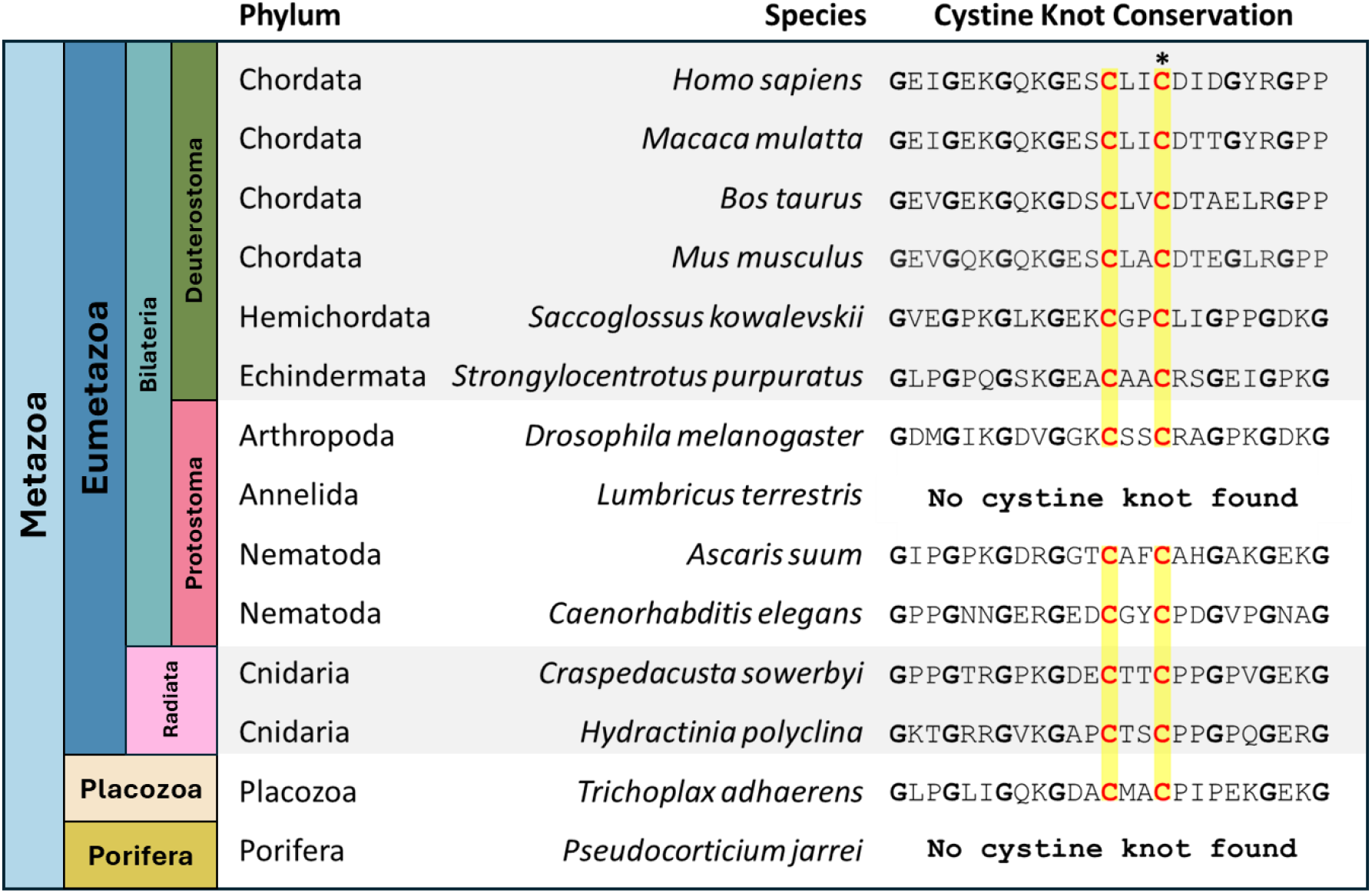
Conservation of cystine knot. Multiple sequence alignment of COL4A1 among 9 metazoan phyla reveals conservation of cysteines allowing for formation of a cystine knot. Cysteines are bolded in red, and the pattern of two cysteine residues is highlighted in yellow. They are absent in the annelidan *L. terrestris* and the poriferan *P. jarrei*. The asterisk indicates the cysteine used to identify the position of the motif in the collagen IV chain.

The cysteines that form the cystine knot were found in all COL4A1 sequences examined from various metazoan phyla, except for the annelidan *L. terrestris* and the poriferan *P. jarrei*, which possess significantly shorter COL4A1 sequences than the other species analyzed. This suggests that a cystine knot may not be essential for stabilizing shorter collagen IV collagenous domains. Our analysis revealed that the cystine knot is typically situated between 907 and 990 amino acids from the N-terminal edge of the NC1 domain, consistently positioned within an interruption in all species that contain it. This implies that the cystine knot is an important evolutionary element in longer interrupted collagenous domains and is likely crucial for their stabilization.

In half (6 out of 12) of the species examined that harbor a cystine knot, this knot contains the first encountered cysteine residue along the canonical folding pathway running from NC1 towards the N terminus (Table 2). However, for longer chains within Bilateria, up to five cysteine residues may precede the cystine knot, suggesting that these collagens may need more cysteines to impart stability. The conservation of the cystine knot in the Placozoan *Trichoplax adhaerens* is particularly noteworthy, as it arose at the onset of the transition into multicellularity. The presence of a cystine knot in this species underscores the evolutionary significance of this conserved cystine knot in providing essential stability to one of the earliest collagen IV structures to arise in evolution. Altogether, this demonstration of conserved cystine knots across Metazoa may suggest a key evolutionary benefit in conferring stability to the collagen IV protomer.

**Table 2.** Cysteines in COL4A1 across species spanning major metazoan phyla.

| Species | # of cysteines | Length from NC1 to *C | # of cysteines before *C |
| --- | --- | --- | --- |
| *Homo sapiens* | 8 | 969 | 0 |
| *Macaca mulatta* | 8 | 907 | 0 |
| *Bos taurus* | 8 | 969 | 0 |
| *Mus musculus* | 7 | 969 | 0 |
| *Saccoglossus kowalevskii* | 8 | 931 | 0 |
| *Strongylocentrotus purpuratus* | 10 | 987 | 1 |
| *Drosophila melanogaster* | 9 | 971 | 2 |
| *Ascaris suum* | 7 | 989 | 0 |
| *Caenorhabditis elegans* | 15 | 990 | 5 |
| *Craspedacusta sowerbyi* | 3 | 972 | 1 |
| *Hydractinia polyclina* | 9 | 966 | 1 |
| *Trichoplax adhaerens* | 3 | 951 | 1 |

## Discussion

In this study, we explored the mechanical, structural, and thermal stability of the collagen IV protomer, which is a foundational component of the BM. By heating and cooling collagen IV protomers, and using AFM to image and analyze the resultant structures, we have been able to identify some key intermediates in the *in-vitro* pathways of unfolding above body temperature and of refolding below body temperature.

Figure 7 summarizes the key structural findings of our study and depicts free energy landscapes at higher and lower temperatures whose features relate to our observations. In response to elevated temperatures (here, temperatures of 37°C, 40°C and 43°C were used in different studies), collagen unfolds. At 37°C (but not 35°C; Fig. S6), we detected localized softening / increased flexibility in a region C-terminal to the cystine knot, which we attribute to localized micro-unfolding of the triple helix. Subsequent thermal fluctuations from this unfolding-prone state can lead to the cooperative unfolding of collagen IV towards its C-terminus. This leads to a metastable intermediate structure with an apparently intact collagenous structure N-terminal to the cystine knot. We detected no other significant intermediate structures on the unfolding pathway, e.g. none that exhibited partial folding of the region between the NC1 domain and the cystine knot. Higher temperatures accelerate the transition to the fully unfolded state, in which all triple-helical structure is lost. This unfolding pathway is consistent with the biphasic thermal denaturation transition observed with bulk CD measurements (Fig. 5D). Upon cooling to room temperature, a collapsed unfolded state (either the intermediate structure or the fully unfolded state) gradually transitions towards the most stable structure, the fully folded collagen IV protomer, likely via the widely accepted zippering mechanism of collagen folding.^7^ However, it encounters barriers along this pathway. Some are due to the ease with which the repetitive G-X-Y sequence can misalign with neighboring α chains and create locally misfolded regions; escaping out of these misfolded states requires unfolding a metastable triple helix to properly align the chains, a process which is notoriously slow and produces a glassy landscape. We observe a broad length distribution of collagenous domains during this refolding process, consistent with the slow and variable dynamics involved in properly zippering up these three chains. Notably, this distribution is not uniform, but is peaked at a length corresponding to folding up to the largest overlapping interruption in the collagen IV sequence. This implies that partially folded states accumulate at this length, where they encounter an entropic barrier to further folding that arises from the necessity to renucleate the triple-helical structure past the interruption. Eventually, many proteins successfully navigate this refolding, but even after overnight incubation at room temperature, approximately ¼ of the proteins remain with a shorter folded contour length than the native ensemble at this temperature.

**Figure 7.**
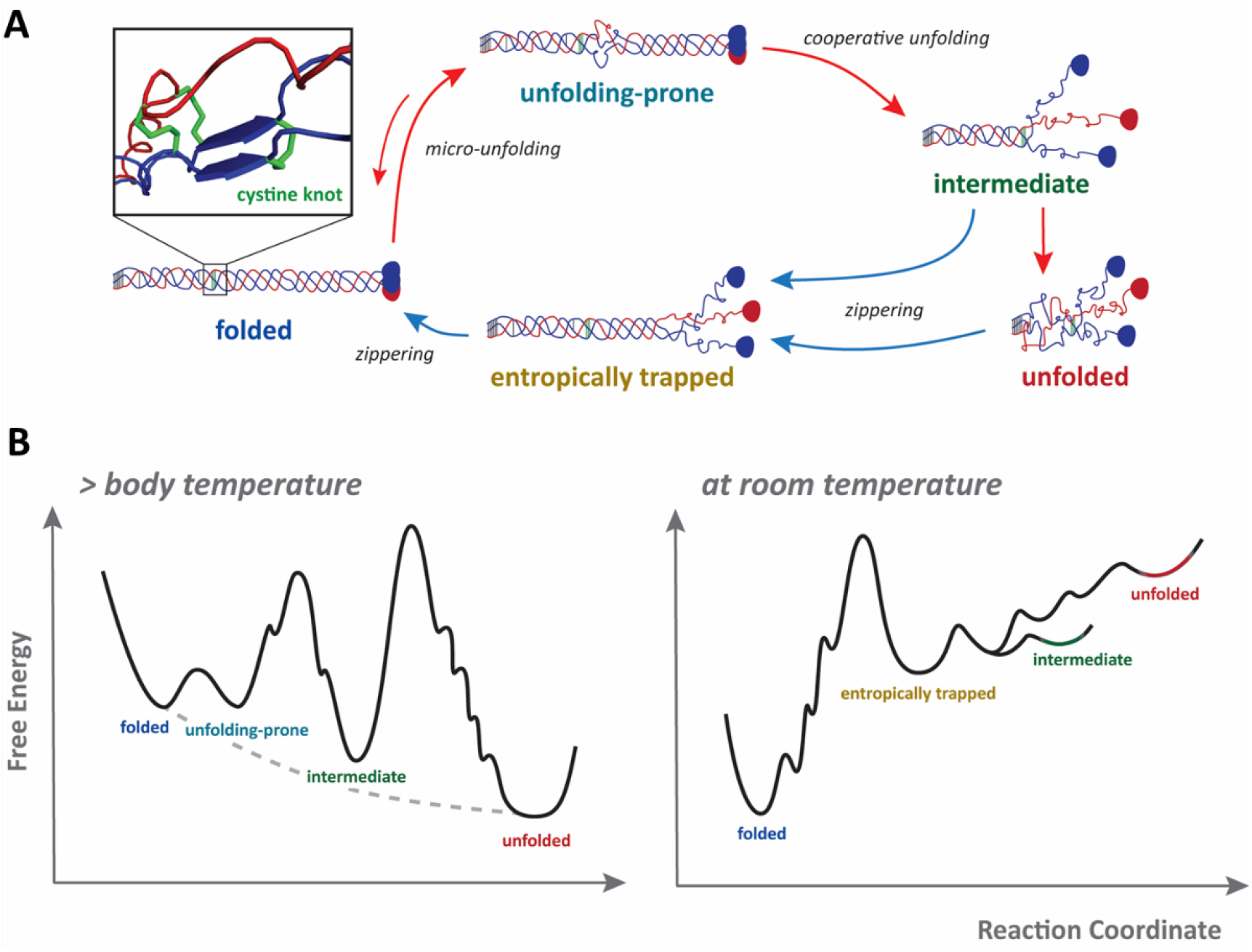
Proposed thermally induced unfolding and refolding pathways. **A**. At body temperature (red arrows), the folded collagen IV protomer exhibits micro-unfolding just C-terminal to the cystine knot. This can lead to the cooperative destabilization of the block C-terminal to the knot, resulting in an intermediate structure in which the chains C-terminal to the cystine knot are unfolded and collapsed into random-coil like structures. Above body temperature, the region N-terminal to the knot can collapse, leading to a fully unfolded state. Upon cooling at room temperature (blue arrows), the intermediate or unfolded state can begin to refold by “zippering” toward the C terminus. Along this pathway, interruptions present local folding barriers, with the largest overlapping interruption presenting an especially high barrier and resulting in an entropically trapped metastable refolding state. Over an extended period, the structure can refold back to native length, albeit with possible chain misalignment within the collagenous domain. Inset shows the predicted structure of the cystine knot, showing interchain disulfide bonds in green, the α1 chains in blue and the α2 chain in red. **B**. Schematic free energy landscapes that are consistent with the observed unfolding and refolding structures. Black lines represent the free energy surfaces for non-reduced collagen IV, while the gray dashed line represents its reduced counterpart. The metastable states observed with AFM imaging are depicted as local minima and labeled accordingly.

Our results demonstrate the crucial role of disulfide bonds, including the cystine knot, in conferring stability to the collagen IV protomer. The clearest demonstration of this is the inability of reduced collagen IV to refold under our experimental conditions. By contrast, non-reduced collagen can refold, albeit very slowly. Consistent with this role of disulfide bonds in promoting refolding, we found that collagen I (lacking cysteine residues) denatures irreversibly, while collagen III (containing a cystine knot at its C-terminus) can refold following thermal denaturation (Fig. S9), as seen in previous studies.^25,43^ Disulfide bonds between α-chains provide two functions promoting refolding: they hold the three α-chains together, significantly facilitating nucleation of refolding (making it a unimolecular rather than trimolecular kinetic process); and their presence maintains local alignment of the three chains, enabling zippering with proper chain registry. Disulfides contribute not just to facilitating refolding but also in preventing unfolding: chemically reduced collagen IV unfolds faster at 37°C than its disulfide-containing counterpart. Thus, the thermal stability collagen IV is enhanced by disulfide bonds.^13^ With both CD spectroscopy and AFM, we observed disulfide-containing collagen IV to unfold in two cooperative steps, in which the intermediate state maintains structure only in the region N-terminal to the cystine knot, presumably because of the many interchain disulfide bonds in this region. Reduced collagen IV unfolds in a single step, albeit less cooperatively (as indicated by the slope of the CD denaturation curve), with the crucial cystine knot no longer present to shield the N-terminal region from destabilization. While our experiments cannot separately evaluate the role of the cystine knot from other more N-terminal disulfides that link the three chains, they do reveal the critical role of the cystine knot as a clamp that prevents complete unfolding of the collagenous domain and that maintains local association and registry to permit more facile refolding of the protein. The importance of these knot-forming two cysteine residues per chain is reinforced by our examination of other collagen IV genes: this pattern of cysteines is evolutionarily conserved throughout Metazoa, consistently located roughly 300 nm from the edge of the NC1 domain. Interestingly, this distance aligns with the average reported contour length for fibrillar collagens, suggesting that collagens longer than 300 nm, particularly those with interruptions, may require interchain disulfides for additional stability.

How does unfolding initiate? Micro-unfolding can occur at shorter (∼5 aa) and longer (∼80 aa) length scales, with the longer domains considered to be a precursor to global unfolding of the collagen triple helix^16^ Thus, we consider three mechanisms by which the collagens of our study can initiate unfolding: end-fraying, micro-unfolding at interruptions, and micro-unfolding of triple-helical regions. End-fraying would be an entropically less costly means to initiate unfolding, but interdomain interactions in the NC1 trimer (*T*_m_ = 66°C) and N-terminal disulfide bonds prevent this from happening in collagen IV.^39,44^ Similarly, the type III collagen we have used includes a N-terminal disulfide-linked propeptide domain and a C-terminal cystine clamp,^22,27^ preventing end-fraying. Only collagen I has unclamped ends; we have managed to capture its end-fraying with our AFM imaging, though this is a rare structure. More insight into the role of internal micro-unfolding comes from examination of contour length changes upon heating. Considering only those collagens that had not yet unfolded, so as to look for signatures of unfolding initiation, we found their mean contour lengths to shorten at 37°C compared to room temperature, by 5% (collagen I), 3.6% (III) and 6.7% (collagen IV). If shortening results from triple helix destabilization, this trend suggests that collagen III exhibits the least destabilization, likely because it has access only to micro-unfolding within triple-helical regions. In contrast, collagen I has the additional option of end fraying, and collagen IV’s interruptions may favor micro-unfolding. While length compression associated with the formation of micro-unfolded bubbles was revealed, our global bending stiffness analysis did not demonstrate the expected softening that we expected to accompany unfolding. Because micro-unfolding is an inherently short structural feature,^16^ we applied our more localized (30 nm; ∼90 aa) sequence-dependent bending analysis to collagen IV. This approach revealed that the region immediately C-terminal to the cystine knot can undergo micro-unfolding, as indicated by increased flexibility in this region at elevated temperatures. The thermal lability of this region makes it highly likely that this site is the nucleation point for thermally induced unfolding of collagen IV.

Folding of the triple helix follows distinct dynamics from the folding of most proteins. Its linear structure forms by “zippering”, which *in vivo* proceeds from the C to the N terminus.^7^ Our results reveal that refolding of collagen IV is also possible in the N- to C-terminal direction. To our knowledge, our study provides the first evidence that this folding direction can be followed by a collagen protein. N-to-C folding directionality is promoted in collagen IV by the aforementioned cystine knot, which positions the chains for zippering from this point towards C terminus. Dölz and Engel previously investigated the potential for collagen IV to undergo N-to-C folding from the cystine knot but found no evidence that this could occur.^12^ Why do we see refolding in this direction while they did not? Most of their studies used collagen IV dimers with an intact NC1 hexamer, which provided a nucleation point for their observed refolding in the canonical C-to-N direction.^13^ To interrogate folding from the cystine knot, they incubated collagen IV in 0.1 M acetic acid, which dissociated the NC1 hexamer. They did not observe C-to-N refolding under these conditions. Could the neutral pH in our study be the reason we observed collagen IV to refold N-to-C? A pH-dependent effect could indicate participation of salt bridges in facilitating refolding.

Since in collagen IV, interruptions can serve as nucleation barriers for propagation of triple helix zippering (such as the largest overlapping interruption as identified in our study), we looked to the sequences flanking these interruptions for insight into the directionality of collagen IV folding. Natural interruptions are often flanked by atypical G-X-Y sequences, which typically contain a higher concentration of charged residues – which could form salt bridges – at their C-terminal edge and an elevated imino acid content, on average, in the two triplets N-terminal to interruptions.^45^ Both types of sequences stabilize triple helix structures at neutral pH; however, at acidic pH, salt bridges will not form. The entropic cost of renucleating a triple helix C-terminal to an interruption is thus not compensated by favorable electrostatic interactions at acidic pH, likely explaining why folding could not propagate in the N-to-C terminal direction in the previous study.^12^ At neutral pH, salt bridge formation provides an electrostatic clamp to stabilize and renucleate the triple helix structure C-terminal to an interruption.^23^ Thus, a salt-bridge-mediated folding mechanism, coupled with the imino-rich region flanking an interruption, may explain how collagen IV can autonomously refold *in vitro*. These special sequences flanking endogenous interruptions may also contribute to the pathology of mutations in collagen that introduce an interruption: they would lack the atypical sequences necessary for efficient folding around the mutated site.

Despite collagen’s inherent instability at body temperature *in vitro*, it is somehow able to fold and maintain stability throughout its physiological journey from synthesis to incorporation into the tissue matrix. We speculate about potential mechanisms that could reconcile these *in vitro* and *in vivo* differences in stability. Heat Shock Protein 47 (HSP47) and SPARC, along with macromolecular crowding effects, can contribute to collagen IV stability *in vivo*. HSP47 maintains properly folded collagen during transit from the ER to the cis-Golgi, binding at near-neutral pH in the ER and dissociating in the ER-Golgi intermediate compartment, demonstrating a regulatory mechanism influenced by the changing chemical environment.^46–48^ Similarly, chaperones like SPARC have been implicated in intracellularly stabilizing collagen IV prior to their potential role in proper BM assembly post-secretion.^46,49^ Despite these mechanisms, ensuring collagen remains properly folded from the ER to its incorporation into the BM is not fully understood. Macromolecular crowding has been shown to promote protein thermodynamic stability *in vitro* by altering solvent properties and enhancing protein-protein interactions.^50^ By contrast, most *in vitro* studies, including our own, use collagen in dilute conditions. For example, our AFM studies used highly diluted protein samples (∼0.2 μg/mL) to ensure we could image non-overlapping proteins. Because of this extremely low concentration (∼10^6^ times less than the physiological environment^51^), collagen’s structure is likely more susceptible to thermal denaturation. The involvement of mechanisms such as chaperone activity and macromolecular crowding suggests that *in vivo* conditions are essential for maintaining collagen’s structural integrity. This highlights the complexity of maintaining collagen stability en route to extracellular assembly, offering insights into how an initially unstable building block can form structurally robust tissues.^52^

In conclusion, this work provides new insights into the complex dynamics of collagen IV, particularly its thermally induced unfolding and refolding mechanisms. The ability of AFM imaging and analysis to capturing and identify individual configurations in the ensemble of molecules allowed us to identify a partially unfolded intermediate state of collagen IV involving the cystine knot, and to identify the location and identity of a major structural barrier to refolding. We elucidated a key role for the cystine knot in facilitating autonomous refolding towards the C terminus, a direction opposite to collagen’s natural folding. The evolutionary conservation of the cystine knot in collagen IV across diverse metazoan phyla highlights its functional importance, as its consistent placement within the collagenous domain is likely essential in stabilizing longer domains. Our observations offer new insights into the mechanisms governing collagen IV’s response to temperature, and highlight its distinct behavior compared to the less structurally complex continuously triple-helical collagens.

## Materials and Methods

### Collagen Purification and Sources

We adapted a previously published protocol to obtain heterotrimeric (α1)_2_α2 PFHR9-derived collagen IV from PFHR-9 medium.^13,32^ A full protocol is provided in reference 53. Briefly, PFHR-9 cells were expanded in 1700-cm^2^ roller bottles (Corning® – CAT# 430852) and maintained at 37°C. The cultured media was harvested every 24-hour period once cells reached confluency. The media used to harvest was DMEM GlutaMax (Gibco™– CAT# 10566016) supplemented with 5% FBS and containing 50 μg/mL ascorbate 6-phosphate (non-hydrolyzable analog of ascorbic acid), and 10 mM potassium iodide. Ascorbic acid serves as an essential cofactor for collagen post-translational-modification enzymes, enhancing collagen secretion.^54^ Potassium iodide inhibits collagen IV cross-linking into dimers, promoting the isolation of fully monomeric protomers by preventing peroxidasin activity on the NC1 domains.^39^

Acetic acid was added to the harvested media to a final concentration of 0.2 M and collagen was precipitated by adding 1.2 M NaCl and allowing the sample to stir at 4°C overnight. The pellet was collected by spinning at 18,500 g at 4°C, then resuspended into 10 mM Tris-HCl, 0.5 M Urea, 25 mM NaCl at pH 7.0. The sample was loaded onto a 5 mL anion-exchange QMA column (Waters™ – CAT#WAT023525), where collagen IV does not bind under these conditions. The flowthrough contained large amounts of collagen IV, and other globular protein contaminants, and was purified further by loading onto a cation-exchange SP FF HiTrap column (Cytiva© – CAT#17515701). Here, most of the contaminants bound to the column, while most of the collagen IV was found in the flowthough. Collagen IV was concentrated in 0.5 M acetic acid by running through an Amicon® Ultra Centrifugal Filter with a 100 kDa MWCO (Millipore® – CAT#UFC8100), and the final sample was stored at 4°C.

Heterotrimeric (α1)_2_α2 rat tail tendon-derived collagen type I was purchased from Trevigen (Cultrex®, 3440-100-01) and is acid-solubilized collagen with a stock concentration of 5 mg/mL in 20 mM acetic acid. Homotrimeric (α1)_3_ bovine pro-N collagen III was extracted from fetal bovine skin following a previously published protocol and was a gift from Yoshihiro Ishikawa.^55^

### Biochemical Preparations

Collagen IV was reduced by mixing an equal volume of the stock (∼0.3 mg/ml in 0.5 M acetic acid) with 0.25 M TCEP-HCl (Sigma-Aldrich®, C4706) at room temperature. After 1.5 hours, the sample was transferred to a pre-wetted dialysis cup (20 kDa MWCO, Thermo Scientific™, 69590) and dialyzed overnight into a large volume of PBS at 4°C. Non-reduced collagen IV was prepared by adding water instead of TCEP-HCl and dialyzing the sample similarly using a dialysis cup.

Samples were run with SDS-PAGE (6% separating gel, 4% stacking gel) to confirm reduction of collagen IV. The samples were prepared for loading by adding 4X protein loading dye, under reducing (includes 20% β-mercaptoethanol) or non-reducing conditions, and boiling for 10 minutes at 80ºC. The gel was run at 220 V for one hour, or until the dye front was 0.5 cm above the bottom edge of the gel. The resolved gel was stained with Coomassie Blue R-250 for 30 minutes, and destained with destaining solution (4% methanol, 1% acetic acid) for one hour.

### Circular Dichroism Spectroscopy

For conducting CD measurements, the stock collagen IV (0.5 M acetic acid) was diluted to a final collagen concentration of ∼0.1 mg/mL in the appropriate solution condition. Spectral scans and variable temperature data were recorded with a JASCO-815 spectrophotometer (UBC Chemistry) using a quartz cuvette with a 0.5 mm path length. The contribution of triple helix structure to the spectra was quantified by *R*_pn_, the ratio of the intensity of the positive peak at 222 nm to the negative peak at 197 nm.^56^ Thermal denaturation curves were obtained by monitoring the mean molar ellipticities (mdeg) at 222 nm. The measurement parameters used are shown in Table S1. The thermal denaturation profiles were plotted by smoothing the data using a window size of 30 as a moving average, with no overlap. The error bars represent the standard deviation within each independent window segment. The curve was fit to a mono- or biphasic sigmoidal function using Igor Pro 8.04.

### Atomic Force Microscopy Imaging

Collagen IV was diluted into PBS (pH 7.0) at the appropriate temperature to a final concentration of ∼0.2 μg/mL. Collagens I and III samples were prepared in 150 mM sodium acetate (pH 5.5), unless otherwise stated. The collagens were allowed to incubate at the given temperature in solution for different time periods before depositing 50 μL of the diluted sample on freshly cleaved mica (Highest Grade V1 AFM MicaQ Discs, 10 mm; Ted Pella, Redding, CA). The samples were left to sit on mica for 10-30 seconds, and excess unbound proteins were removed by rinsing with ultrapure water at the appropriate temperature, followed by drying the mica using filtered air. All protein imaging took place under dry conditions, and the solution conditions and temperature of the samples refer to the conditions at the time of deposition onto the mica surface. To ensure thermal equilibration, solutions along with other materials including pipette tips, mica disks, and water were subjected to overnight incubation at the desired temperature prior to use. Imaging was done with an Asylum Research Jupiter XR AFM (access graciously provided by Emily Cranston, UBC Engineering and Forestry) using AC tapping mode in air. AFM tips with a 160-kHz resonance frequency and 5 N/m force constant (MikroMasch, HQ: NSC14/AL BS; Sofia, Bulgaria) were used.

### Chain Tracing and Analysis

Our custom MATLAB code SmarTrace was used to determine the persistence length of collagen.^57^ The code was validated extensively in previous work on collagen.^8,58,59^ Chains were traced to subpixel resolution starting from the internal edge of the C-terminal NC1 domain. The analysis of a global persistence length assumed uniform flexibility along the chains. Persistence length was determined from the dependence of tangent vector correlation and mean-squared end-to-end distance on the segment length:^28,60^

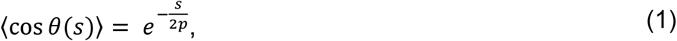

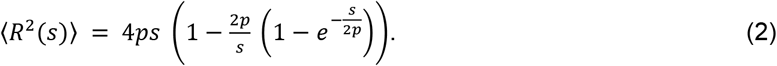

To study the sequence dependence of collagen’s flexibility, the local persistence length was determined as a function of position along the collagen contour.^8^ Briefly, the analysis determined the effective persistence length of a collagen chain segment of length Δ*s* = 30 nm, with outcome associated with the position *s* at the centre of this window. To do so, the angular variance was measured for a segment length Δ*s* centered at position *s* from many traced collagen chains; this variance is inversely proportional to the effective persistence length via

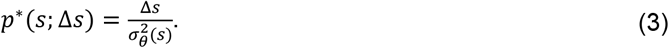

95% confidence intervals were calculated for *p*\*(*s*) using the cumulative probability function of the scaled inverse chi-square distribution.^8^

Because persistence length is an inherently temperature-dependent parameter (more thermal energy leads to enhanced bending fluctuations), determining whether collagen’s structure changes with temperature requires the use of a different parameter. Bending stiffness is used because it is a temperature-independent property, providing a better measure of intrinsic material properties. Persistence lengths obtained from equations (1-3) were converted to bending stiffness values for all subsequent analyses using the following relation:^61^

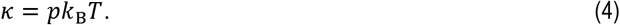

### Multiple Sequence Alignment

Collagen IV sequences from different metazoan species were taken from NCBI. All sequences belong to the collagen IV α1-like subfamily of chains, except for *Ascaris* (α2 chain). Two cysteine residues spaced two residues apart within an interruption were identified in most sequences and subsequently aligned. The NCBI GenBank accession numbers are listed in Table 1.

### Predicted structure of the cystine knot

AlphaFold3was used to predict the local disulfide-bridged structure around the cystine knot.^62^ (G-P-O)_4_ sequences were added to flank the interruption sequences in α1 and α2 that form the cystine knot, to aid in aligning the chains for prediction of the local structure.

## Acknowledgments

We are extremely grateful to Emily Cranston and team from UBC Faculty of Forestry for allowing us to use their Asylum Jupiter AFM (instrument funded by the Canadian Foundation for Innovation). We gratefully acknowledge Yoshihiro Ishikawa for the gift of collagen III, and thank Ben Herring at UBC for access to the Department of Chemistry CD spectrometer. We thank members of the Forde lab for insightful suggestions on this work. We also thank David Sivak and Eldon Emberly for critical feedback on a draft of this manuscript.

This work was funded by the Natural Sciences and Engineering Research Council of Canada (NSERC) through a Discovery Grant to NRF and a Post-Graduate Scholarship, Doctoral (PGS-D) to AAS.

