## Supporting information for "Decoding Collagen’s Thermally Induced Unfolding and Refolding Pathways"

**This PDF file includes:**

Figures S1 to S12
Table S1
SI Reference

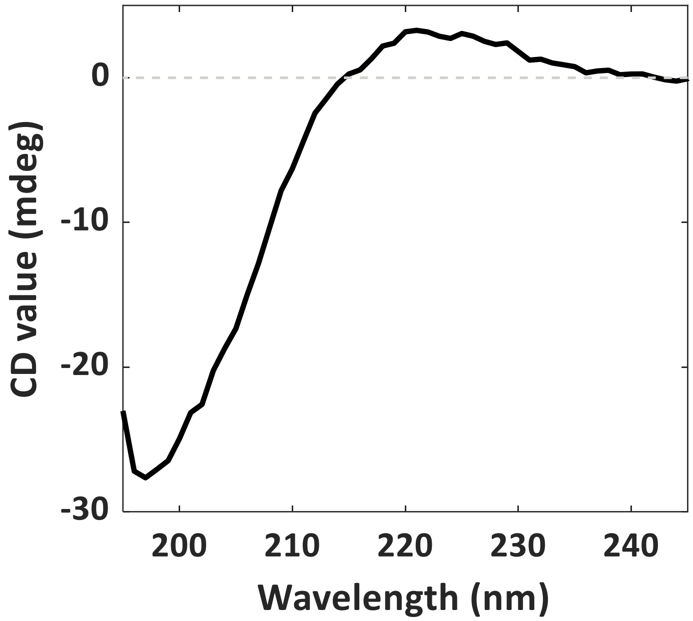


**Fig. S1.** **Circular dichroism spectrum of collagen IV**

The spectrum was recorded of collagen IV in PBS at ambient temperature. It exhibits a maximum at 222 nm and a minimum at 198 nm, with an $R_{pn}$ value of 0.13, indicating triple-helical structure.

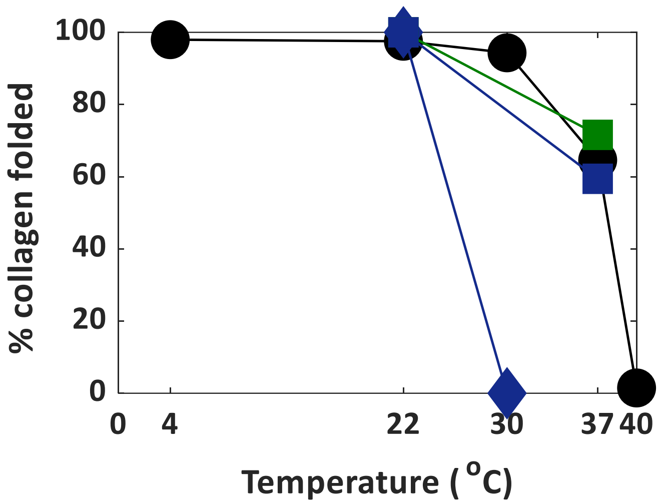


**Fig. S2.** **Temperature-dependent loss of folded collagen**

Collagen was incubated at different temperatures for 30 minutes prior to imaging with AFM. Collagen IV (black circles) was imaged in PBS (pH 7.0). Collagen I was imaged in 150 mM sodium acetate buffer, pH 5.5 (blue squares) and in 100 mM KCl, 1 mM HCl, pH 3.0 (blue diamonds), while collagen III (green squares) was imaged in 150 mM sodium acetate buffer, pH 5.5. Lines are to guide the eye.

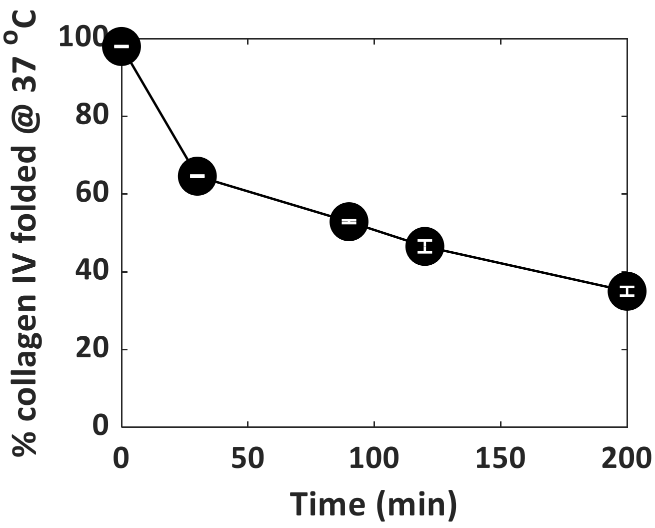


**Fig. S3.** **Time-dependent loss of folded collagen IV**

Collagen IV was heated in PBS at 37°C for different time periods followed by imaging with AFM.

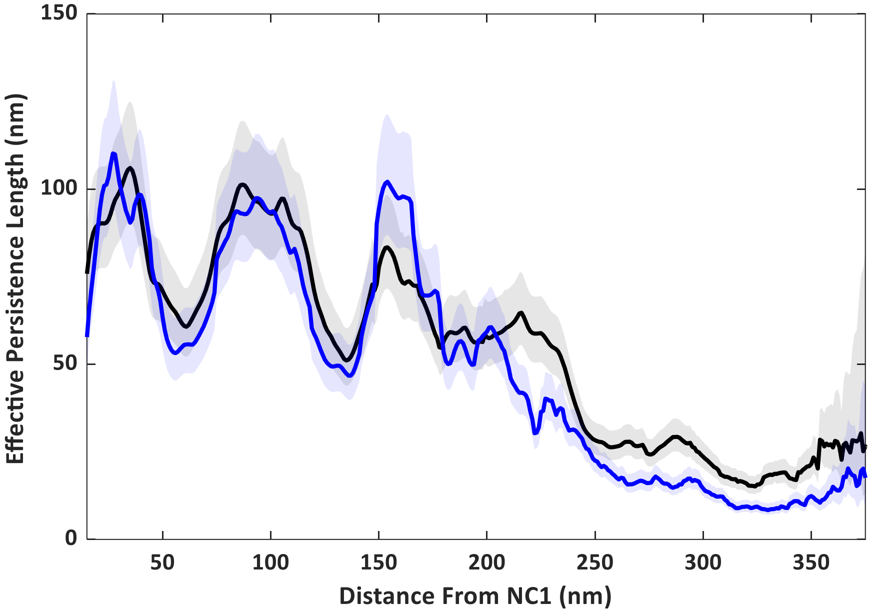


**Fig. S4. Effective persistence length of tissue-derived vs PFHR9-derived collagen IV**

PFHR9-derived (black, *N*=305) and tissue-derived collagen IV (blue, *N*=262, reference (1)) were imaged in 100 mM KCl with 1 mM HCl at pH 3.0.

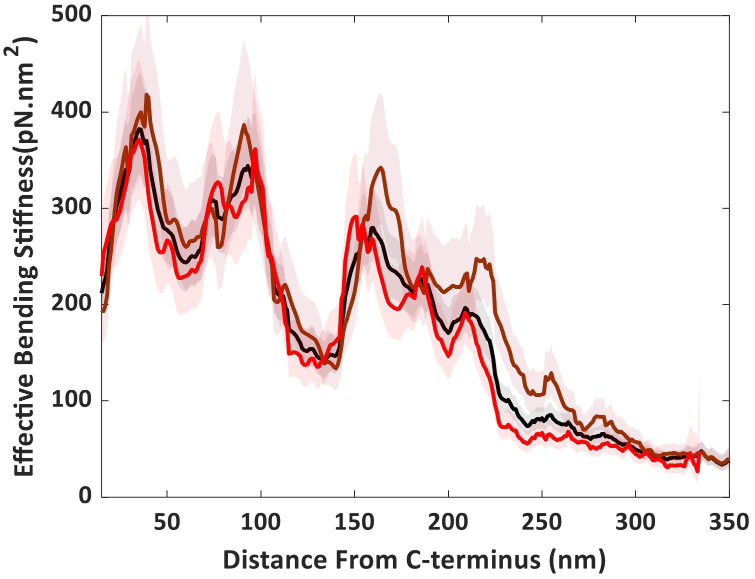


**Fig. S5.** **Effect of contour lengths on the flexibility profile of collagen IV at 37°C**

Effective bending stiffness profile of collagen IV deposited after a 30-minute incubation at 37°C (black; *N*=413, same as bright red in Fig. 2 of the main manuscript). The resulting profile when imposing a contour length selection for only those chains >350 nm (dark red, *N*=202) or <350 nm (red, *N*=211).

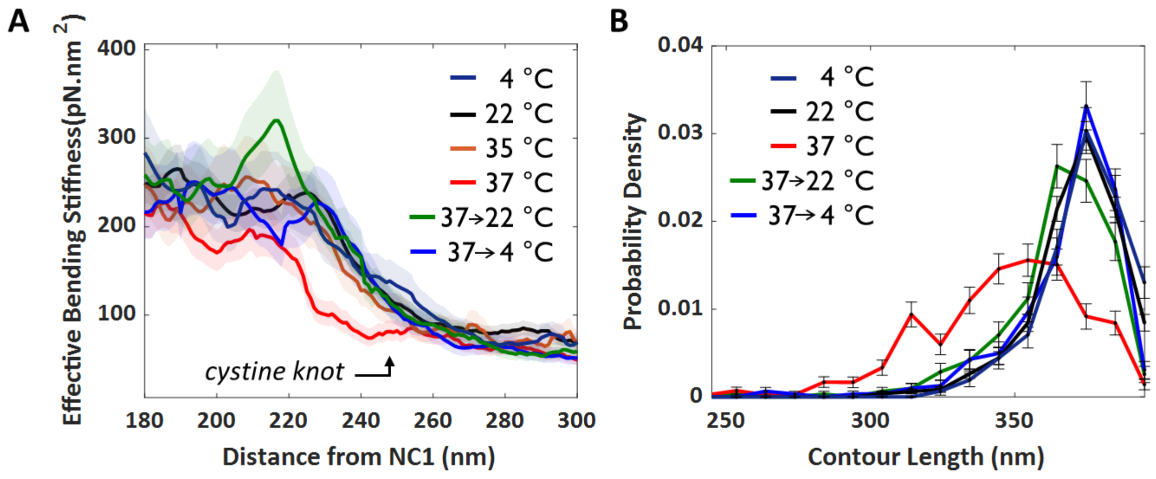


**Fig. S6.** **Restoration of the initial destabilized region and contour length upon cooling**

**A.** Effective bending stiffness profiles near the putative cystine knot at different temperatures, including upon cooling from 37°C to overnight at room temperature or at 4°C. Structural destabilization at this site was not observed at 35°C, just below body temperature. **B.** Distribution of contour lengths of collagen IV with visibly intact NC1 domains, at different temperatures, including upon cooling from 37°C to room temperature and to 4°C.

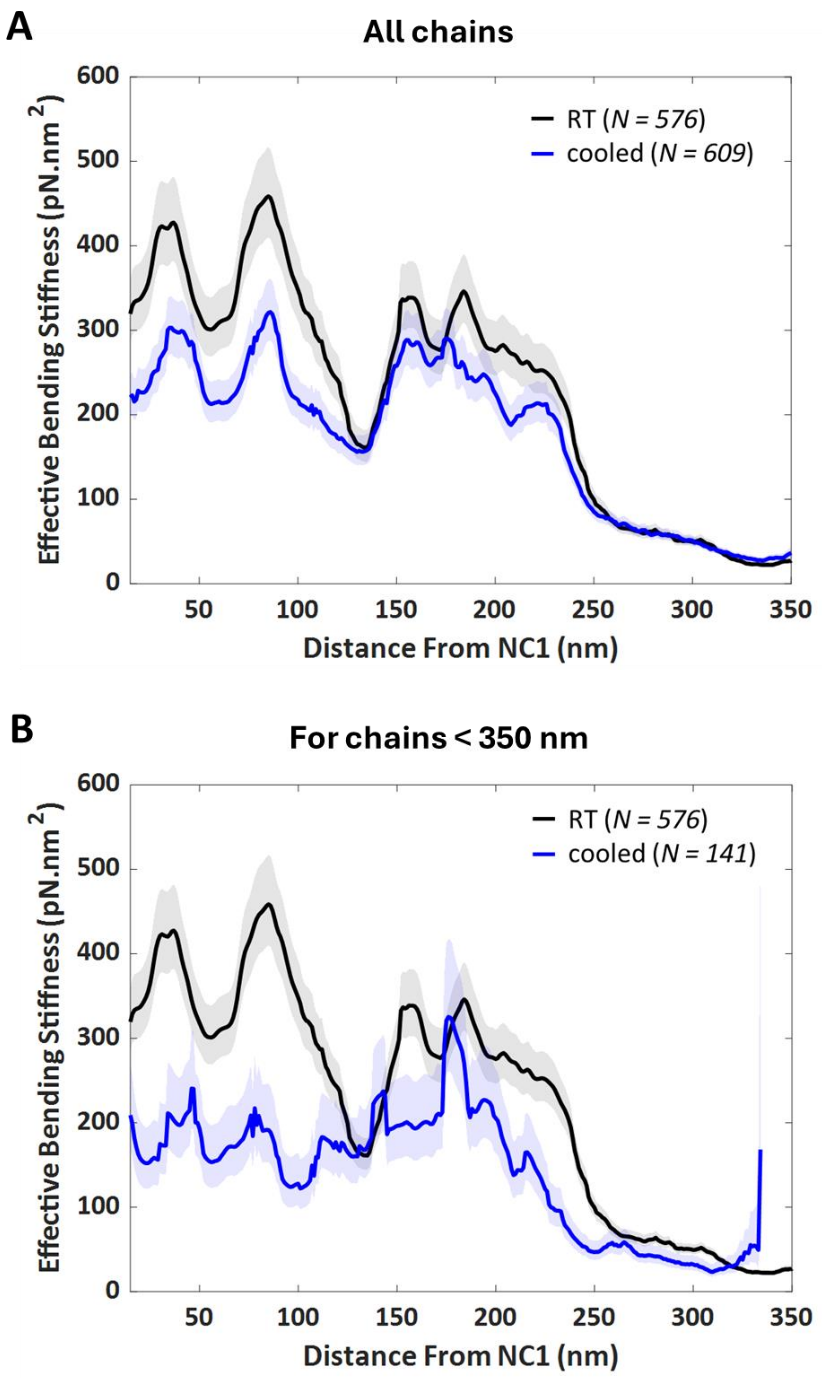


**Fig. S7.** **Effective bending stiffness profile of refolded collagen IV**

Related to Figure 4 in the main manuscript. **A.** Profile including all traced chains from the refolded sample (blue), compared to the native profile at room temperature (black). **B.** Resultant profile if only chains <350 nm in contour length (blue) are included in the analysis.

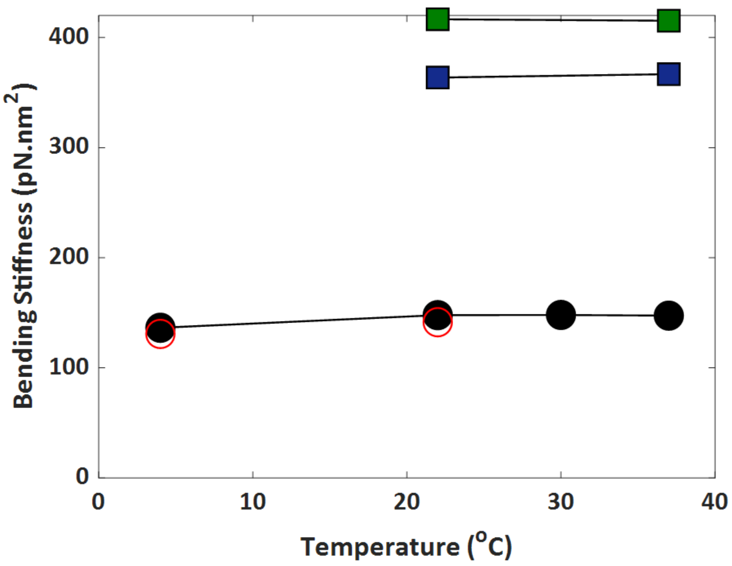


**Fig. S8.** **Global bending stiffness of collagens as a function of temperature**

Collagen was plated in PBS (circles) or 150 mM sodium acetate (squares) at different temperatures. The average value of persistence length resulting from analysis with equations (1) and (2) was used in equation (4) to determine the global bending stiffness of each collagen type at each temperature. Collagen III (green) is stiffer than collagen I (blue) and collagen IV (black). Collagen IV samples that were heated at 37°C for 30 minutes followed by overnight cooling at 4°C or at room temperature are shown with red circles.

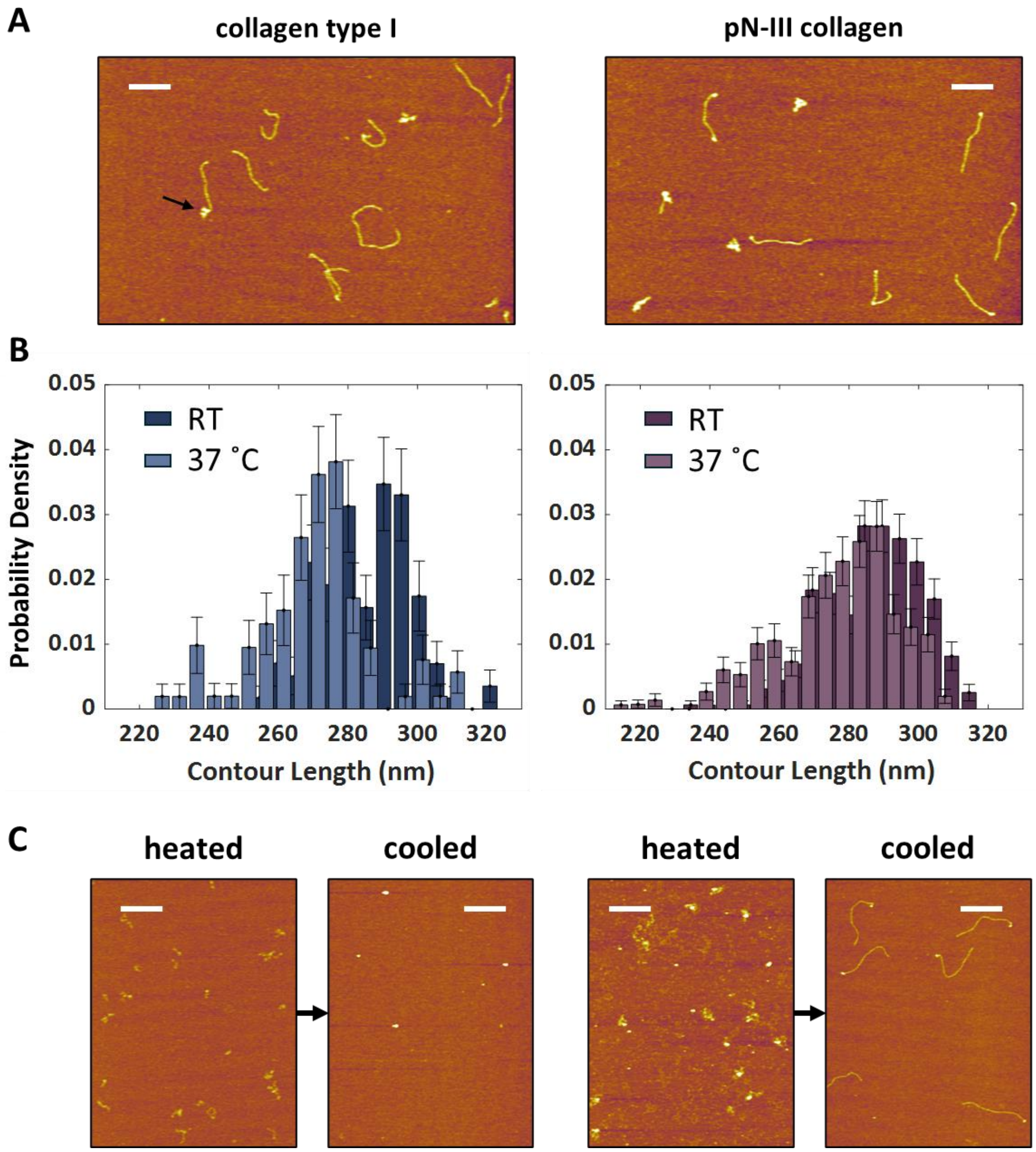


**Fig. S9.** **Fibrillar collagen denaturation and refolding**

**A.** AFM images of collagens I and III after heating at 40°C for 30 minutes. Black arrow identifies an example of end-fraying. Scale bars = 200 nm. **B.** Probability distributions of the contour lengths at room temperature and 37°C for collagen I (left) and pN-III (right). **C.** AFM images of heated collagen I (left) and collagen III (right) at 40°C for 30 minutes followed by cooling overnight at room temperature. Scale bars = 200 nm.

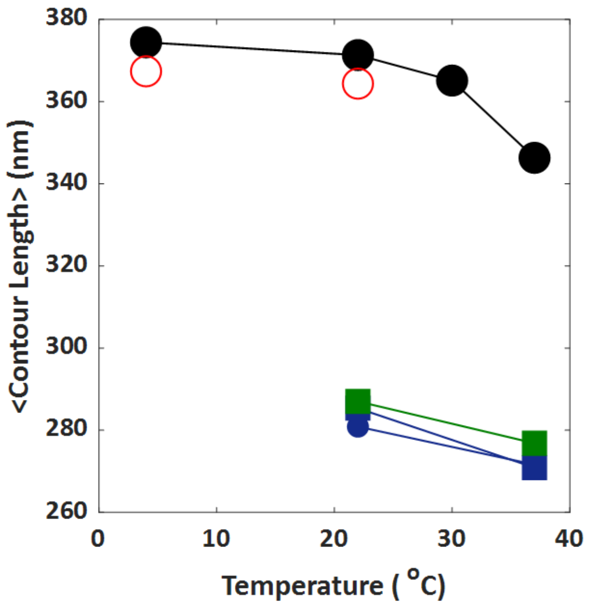


**Fig. S10.** **Contour length of collagens as a function of temperature**

Collagen was plated in PBS (circles) or 150 mM sodium acetate (squares) at different temperatures. The average contour length decreases with temperature for intact collagens I (blue), III (green), and IV (black). Collagen IV samples that were heated at 37 °C for 30 minutes followed by overnight cooling at 4 °C or at room temperature are shown with red circles. Collagen IV has a longer contour length than the fibrillar collagens.

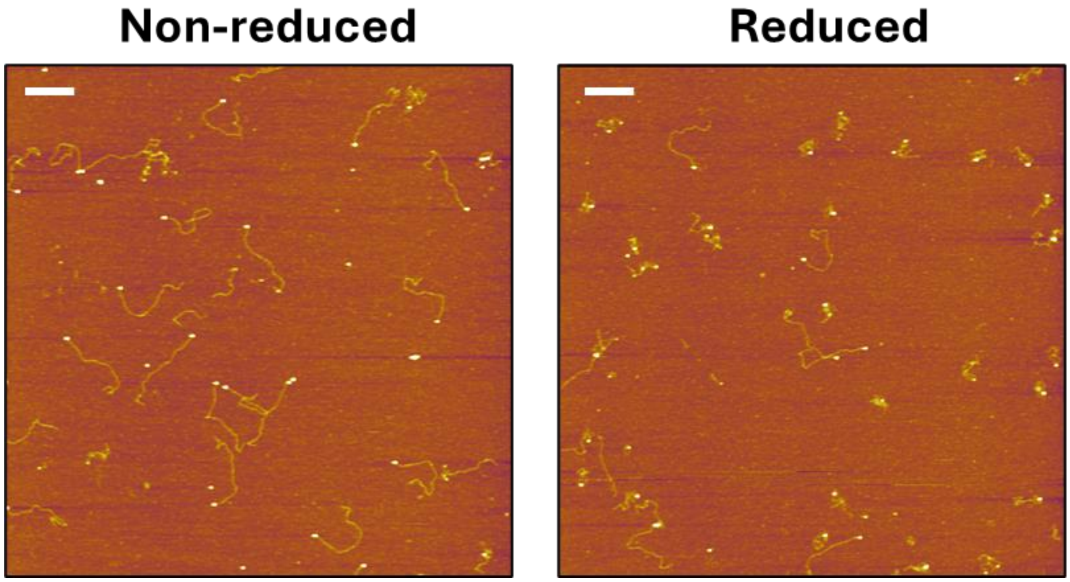


**Fig. S11.** **Effect of reduction on the thermal stability of collagen IV**

Collagen IV was reduced by treating with 0.25 M TCEP-HCl for 30 minutes. Non-reduced and reduced collagen IV were heated at 37°C for 30 minutes in PBS followed by imaging with AFM. Scale bars = 200 nm.

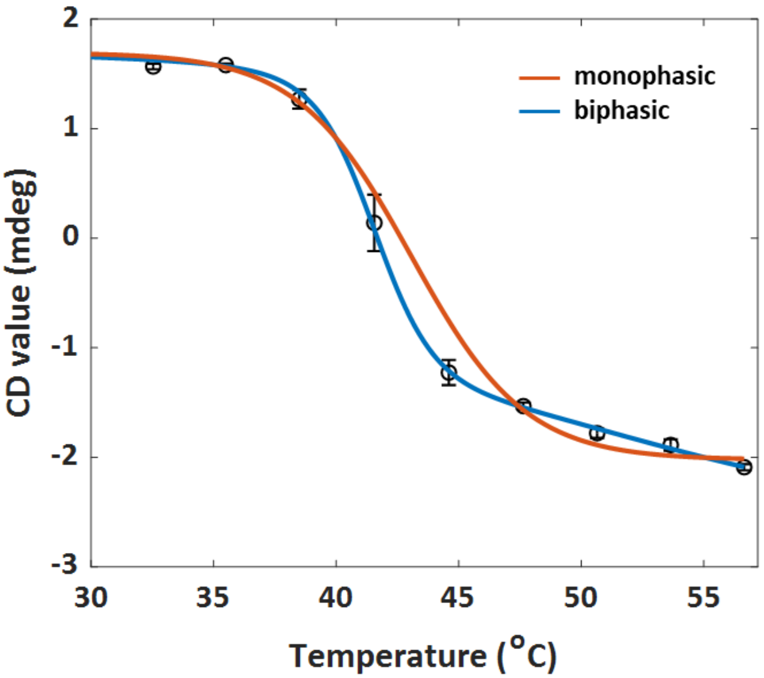


**Fig. S12.** **Melting profile of non-reduced collagen IV in acidic conditions.**

Thermal denaturation profile was obtained by monitoring the CD signal at 222 nm. Non-reduced collagen IV in 0.1 M acetic acid is well described by a biphasic transition. For the monophasic fit, the first transition is at 43.1 ± 0.2 °C with a slope of 0.43 ± 0.01 mdeg/°C and a $\chi_r^2 = 6$. For the biphasic fit, the first transition is at 41.6 ± 0.4 °C with a slope of 0.85 ± 0.19 mdeg/°C and the second transition is at 51.3 ± 6.8 °C with a slope of 0.16 ± 0.06 mdeg/°C and a $\chi_r^2 = 2$.

**Table S1.** **Circular Dichroism parameters.**

| Spectral Scan | Variable Temperature |
|---|---|
| Sensitivity: standard | Wavelength: 222 nm |
| D.I.T (integration time): 1 sec | D.I.T (integration time): 1 sec |
| Bandwidth: 1 nm | Bandwidth: 2 nm |
| Scanning speed: 100 nm/min | Ramp rate: 0.5°C/min |
| Start: 250 nm | Start: 30°C |
| End: 190 nm | End: 60°C |
| Data pitch: 0.2 nm | |
| Accumulation: 5 | |
